# Ovarian germline stem cell dedifferentiation is cytoneme dependent

**DOI:** 10.1101/2024.11.19.624350

**Authors:** Catherine Sutcliffe, Nabarun Nandy, Raluca Revici, Heather Johnson, Shukry J. Habib, Hilary L. Ashe, Scott G. Wilcockson

**Affiliations:** Faculty of Biology, Medicine and Health, University of Manchester, Manchester M13 9PT, UK; Department of Biomedical Sciences, Ru du Bugnon 7, 1005 Lausanne, Switzerland; Developmental Signalling Laboratory, The Francis Crick Institute, London NW1 1AT, UK

**Keywords:** Germline stem cell, *Drosophila* ovary, cytoneme, dedifferentiation, maintenance, BMP signalling, Enabled, S2 cells, BMP beads

## Abstract

Progenitor cell dedifferentiation is important for stem cell maintenance during tissue repair and age-related stem cell decline. Here, we use the *Drosophila* ovary as a model to study the role of cytonemes in BMP signalling-directed germline stem cell (GSC) maintenance and dedifferentiation of germ cells to GSCs. We provide evidence that differentiating germ cell cysts extend longer cytonemes that are more polarised towards the niche during dedifferentiation to reactivate BMP signalling. The presence of additional somatic cells in the niche is associated with a failure of germ cell dedifferentiation, consistent with the formation of a physical barrier to cytoneme-niche contact and outcompetition of germ cells for BMP. Using BMP beads *in vitro*, we show that these are sufficient to induce cytoneme-dependent contacts in *Drosophila* tissue culture cells. We demonstrate that the Enabled (Ena) actin polymerase is localised to the tips of germ cell cytonemes and is necessary for robust cytoneme formation, as its mislocalisation reduces the frequency, length and directionality of cytonemes. During homeostasis, specifically perturbing cytoneme function through Ena mislocalisation impairs GSC fitness by reducing GSC BMP signalling and niche occupancy. Disrupting cytonemes by targeting Ena during dedifferentiation reduces germ cell BMP responsiveness and the ability of differentiating cysts to dedifferentiate. Overall, our results provide evidence that cytonemes play a fundamental role in establishing polarised signalling and niche occupancy during stem cell maintenance and dedifferentiation.

**Significance Statement:** Fertility depends not only on germline stem cell (GSC) maintenance during homeostasis, but also on the ability of germ cells to reverse their developmental programme and dedifferentiate during ageing- or stress-induced GSC loss. Here, using the *Drosophila* ovary as a model, we show that cytonemes are required for GSC asymmetric signalling, niche occupancy and fitness during homeostasis. During dedifferentiation, we demonstrate that cytonemes are necessary for differentiating germ cells to re-access the self-renewal signal and establish polarised signalling, recolonise an empty niche and dedifferentiate to germline stem cells. Our data identify cytonemes as a novel target for strategies aimed at improving stem cell fitness or dedifferentiation, which has huge therapeutic potential for the regeneration of damaged tissues.

## Introduction

Stem cells reside in a niche, a specialised microenvironment that provides physical support and signals to control stem cell maintenance (1). Actin-rich signalling filopodia called cytonemes (2, 3) mediate stem cell-niche communication in different contexts. For example, *Drosophila* haematopoetic niche cell cytonemes deliver Hedgehog (Hh) signals to blood progenitor cells to maintain their fate. Cytonemes are disrupted upon infection, resulting in ectopic differentiation of progenitors into hemocytes to strengthen the immune response (4). *Drosophila* adult muscle progenitors (AMPs) extend cytonemes loaded with FGF receptors to adhere to wing disc niche cells and receive the FGF signal, resulting in asymmetric signalling and AMP-niche organisation. There are two spatially restricted niches: each expresses a different FGF ligand and supports a distinct population of muscle-specific AMPs due to specific cytoneme-niche contacts (5). Embryonic stem cells (ESCs) extend cytonemes carrying Wnt receptors to pair with co-cultured trophoblast stem cells (TSCs), receive the self-renewal Wnt signal and initiate formation of embryo-like structures (6). In contrast, the cytonemes from ESCs that have transitioned to a primed state are unable to form stable interactions with Wnt sources, impairing TSC pairing and synthetic embryogenesis (7).

*Drosophila* ovarian germline stem cells (GSCs) also use cytonemes for short-range niche communication. The GSC niche is composed of specialised somatic cells, including cap cells (CpCs) and escort cells (ECs), which provide self-renewal signals to support 2-3 GSCs per germarium (8, 9). GSCs undergo asymmetric division to form one GSC, which remains in the niche, and one daughter cystoblast (CB) which leaves the niche and differentiates. The key signal promoting stem cell identity is Bone Morphogenic Protein (BMP) signalling, driven primarily through the ligands Decapentaplegic (Dpp) and Glass Bottom Boat (Gbb), resulting in phosphorylation of the transcription factor Mad (10, 11). In addition, GSCs are anchored to the CpCs through E-Cadherin (Ecad) mediated cell adhesion at adherens junctions (12). GSCs are therefore identifiable by their anterior localisation, high pMad levels, and the presence of the spectrosome, a germline-specific spectrin-rich endomembrane organelle that becomes a branched structure (fusome) in more developed cysts (Figure 1A). BMP signalling silences transcription of the differentiation master gene, *bag of marbles* (*bam*) (11, 13). Upon GSC division, the daughter cell that exits the niche has reduced pMad and upregulates *bam* expression, triggering germline differentiation to a CB (14, 15). A single CB undergoes four rounds of mitosis with incomplete cytokinesis to form 2-, 4-, 8- and 16-cell cysts (cc) with interconnected fusomes (16, 17).

**Figure 1:**
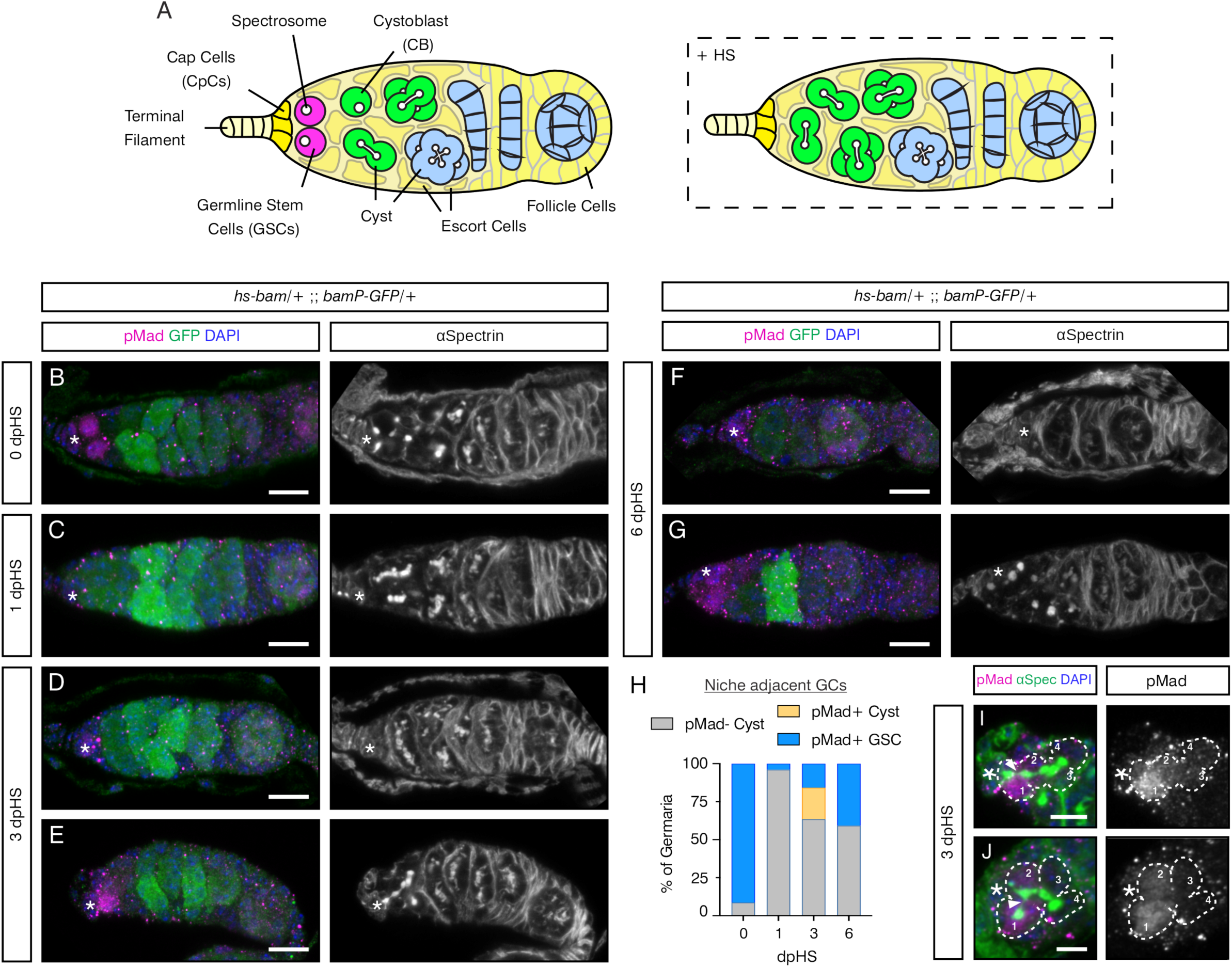
GCs dedifferentiate over time to reoccupy the niche. (A) Schematic of the germarium with the different cell types, as labelled, anterior is to the left. The spectrosome and fusomes are shown in white. The boxed germarium shows post-HS induced *bam* expression. (B-G) Germaria carrying *bam* promoter-GFP and *hs-bam* transgenes immunostained with anti- pMad (magenta), anti-GFP (green), anti-αSpectrin (grey, right panel) and DAPI (blue). Representative germaria are shown for day 0 (B) and 1 dpHS (C), 3 dpHS (D-E) and 6 dpHS (F-G). (*) Cap cells. Scale bar, 10μm. (H) Quantitation of the number of germaria where the GCs contacting the niche are pMad- positive or -negative differentiating cysts or pMad-positive GSCs on 0, 1, 3 and 6 dpHS. The number of germaria counted per time point were 104, 100, 101 and 103 for 0, 1, 3 and 6 dpHS, respectively. (I and J) pMad-positive cysts at 3 dpHS immunostained with anti-pMad (magenta, grey right panel), anti-αSpectrin (green) and DAPI (blue). The image in (I) is from the germarium shown in (E). Connected CBs are outlined and numbered 1 to 4 in the 4-cell cyst. (*) Cap cells, arrowhead shows a fusome undergoing scission. Scale bar, 5μm.

Restricted access to BMP is an important regulatory step that limits stem cell identity to those cells in direct contact with the niche (18). GSC-niche signalling is mediated in part by cytonemes (19). It has been shown that early germ cells (GCs) form actin-rich projections and GSCs extend cytonemes to probe the niche and access Dpp bound by the heparan sulphate proteoglycan Division abnormally delayed (Dally) to promote BMP signal transduction. Dpp signalling induces the formation of more stable microtubule-enriched cytocensors, which also receive the BMP signal but provide feedback inhibition through signal suppression (19). Short cytonemes mediating Hh distribution have also been detected in the germline niche emanating from Hh-producing CpCs toward receiving ECs (20). Thus, cytonemes play an integral role in cell communication in the ovarian germline.

Many factors, particularly those associated with the actin cytoskeleton, have been identified as regulators of cytoneme formation (3). Enabled (Ena), an actin polymerase, is associated with filopodia extension in many cellular contexts (21–24). Ena localises to the tips of lamellipodia and filopodia (24–27), where it recruits Profilin/actin complexes to enhance F-actin polymerization (28, 29) and filopodia formation (30, 31). In addition, Ena localises to focal adhesions and adherens junctions (32). In the latter, Ena plays a role in Ecad localisation (33) and in cytoskeletal regulation at Ecad contact sites (32). Ena is therefore an attractive candidate that could regulate GSC cytoneme formation and niche interaction.

It has previously been shown that transient induction of *bam* expression using a heat shock (HS)-inducible transgene, *hs-bam*, causes complete loss of GSCs from the niche through Bam-induced differentiation within 24 hrs (34). Following this, over the course of a few days, GCs can recolonise the niche and fully recover GSC loss. This is driven by cyst dedifferentiation, resulting in gradual cyst breakdown to form single cells (35). Here, we used this system to test the role of cytonemes in mediating BMP-dependent dedifferentiation. We show that Ena loss-of-function decreases the robustness of cytoneme formation. This reduces GC responsiveness to BMP signalling and niche occupancy, which impairs GSC fitness during homeostasis and precludes niche recolonisation during GC dedifferentiation.

## Results

### Early cysts can dedifferentiate to recolonise an empty GSC niche

To investigate a role for cytonemes in GSC dedifferentiation, we first established the timeline of BMP responsiveness and dedifferentiation using a previously described assay (35, 36). We transiently induced expression of the differentiation factor *bam* throughout the germarium using a *hs-bam* transgene. The number of pMad-positive early GCs was visualised in the days post-HS (dpHS) to quantify the level of dedifferentiation and recovery of niche-associated GSCs. The presence of pMad indicates that the cells are responding to niche derived Dpp signalling. A transgene with the *bam* promoter driving GFP (*bamP-GFP*) (13) was also used to mark Bam-positive differentiating cells. Pre-HS, pMad-positive GSCs are identified by the presence of an anteriorly localised spectrosome (visualised by αSpectrin staining) associated with niche CpCs (Figure 1A, B). GFP is detected in differentiating CBs and developing cysts located more posteriorly (Figure 1B).

One dpHS, pMad- and spectrosome-positive GSCs are absent and the cells directly adjacent to the niche are GFP-positive with branched fusomes (Figure 1C). This indicates that GSCs and early CBs have been lost due to differentiation (Figure 1A). Indeed, almost all germaria contain pMad-negative cysts (Figure 1H; 1 dpHS cf day 0 pre-HS). At 3 dpHS, while ∼60% of germaria still only contain differentiated cells (Figure 1D), ∼40% now contain pMad-positive and GFP-negative cells adjacent to the niche, consistent with dedifferentiation (Figure 1E). pMad accumulation is either associated with a spectrosome-containing GSC in direct contact with the niche, or a cyst containing a branched fusome (Figure 1H). Within these pMad- positive cysts (typically at the 4cc stage) only the cells in direct contact with the CpCs, the source of Dpp, are pMad-positive (Figure 1I, J). Fusome thinning and scission, which are associated with cyst breakdown during dedifferentiation (35), are also evident at the 4 cc stage.

By 6 dpHS, the majority of germaria continue to display a lack of dedifferentiation such that all early GCs are lost, and only later stage cysts are present (Figure 1F, H). This suggests that after 3 dpHS there is no further dedifferentiation of cysts. However, in germaria where dedifferentiation has occurred, pMad- and spectrosome-positive GSCs are detected in the niche (Figure 1G). The proportion of germaria with pMad-positive GSCs that have dedifferentiated is highest at 6 dpHS (Figure 1H). Together, these data show that this post-HS time course can capture different events during GC dedifferentiation, including the ability of cysts to respond to Dpp signalling, followed by cyst breakdown and recolonisation of the GSC niche.

### Dedifferentiating GCs extend cytonemes during niche recolonisation

As we previously showed that cytonemes enable GSCs to access niche Dpp (19), we used the dedifferentiation assay to test whether cytonemes play a role in the dedifferentiation of early cysts by enabling access to niche derived Dpp. To visualise cytonemes during dedifferentiation, we used the germline-specific *nosGAL4VP16* driver to activate a *UASp-LifeAct-GFP* reporter and observed GCs 3 and 4 dpHS. Pre-HS control germaria were also studied, which contain GSCs with niche directed cytonemes (Figure 2A).

**Figure 2:**
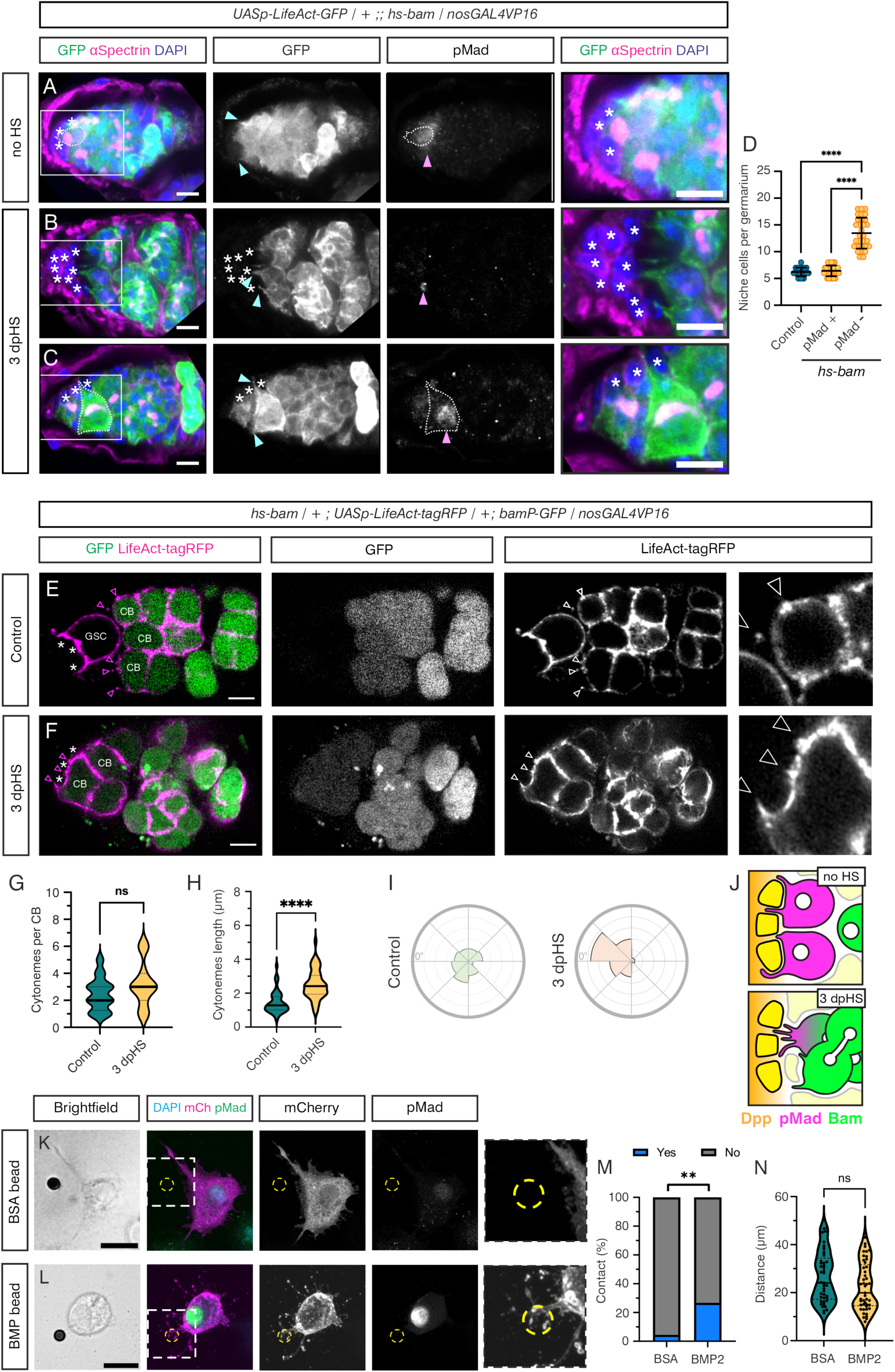
Dedifferentiating GCs extend cytonemes during niche recolonisation. (A-C) Germaria carrying *UASp-LifeAct-GFP* driven by *nosGAL4VP16* and a *hs-bam* transgene immunostained with anti-GFP (green and inset), anti-αSpectrin (magenta), anti-pMad (inset) and DAPI (blue). non-HS control (A), 3 dpHS germaria (B-C) with (B) a pMad negative 8cc extending cytonemes to the niche and (C) a pMad-positive GSC with cytonemes contacting the niche. (*) Niche cells including extra somatic cells, cyan arrowheads mark cytonemes, magenta arrowheads indicate pMad-positive cells, white dotted lines outline pMad positive GSCs. Scale bar, 5μm. (D) Graph shows quantitation of the number of niche cells in *hs-bam* germaria (3 dpHS) that contain pMad-positive dedifferentiating GCs or only pMad-negative GCs, compared to control germaria (HS, with no *hs-bam* transgene). n = 30, data shown as mean ± SD, ANOVA p =<0.001 (control vs pMad- GCs), p= <0.001 (pMad+ GCs vs pMad- GCs) (E, F) Images of live germaria carrying *UASp-LifeAct-tagRFP* driven by *nosGAL4VP16* (magenta and inset), a *bamP-GFP* reporter to mark differentiating cells (green and inset) and a *hs-bam* transgene. non-HS control (E) and 3 dpHS germaria (F) show the cytonemes on a GSC and CBs (E, arrowheads) or a CB occupying the niche (F, arrowheads). Higher magnification views of *LifeAct-tagRFP* labelled CB cytonemes are shown in the right panel. (*) Niche cells. Scale bar, 5μm. (G-I) Quantification of the number of cytonemes per CB (G), cytoneme length (H) and orientation relative to the niche (0°), each interval is 10% (I). n=50 (*hs-bam*) and 46 (control), Welch’s unpaired two-tailed t test, p <0.0001, ns not significant. (J) Schematic of the germarium niche showing no HS GSCs and a 3 dpHS cyst which extends cytonemes toward the niche and is BMP responsive. (K, L) S2R+ cells transfected with Tkv-mCherry and FLAG-Mad expression plasmids and incubated with BSA (K) and BMP2 (L) coated beads. S2R+ cells were immunostained with anti-tdTomato (magenta and inset) and anti-pMad (green and inset). Bead locations are marked with yellow dashed lines. Brightfield images and higher magnification views of the Tkv-mCherry channel are shown. Scale bar, 10 μm. (M) Quantification of the contacts between BSA and BMP2 coated beads and S2R+ cells (examples are shown in K, L). n = 45, Fisher’s exact test, p = 0.0071. (N) Quantification of the distance between the bead and the centre of the cell nucleus. Mann-Whitney p = 0.0781.

Post-HS, germaria in various stages of dedifferentiation were identified. Some germaria have only early or late cysts remaining and lack cytonemes (Figure S1A, B). In contrast, other germaria show pMad-negative early cyst cells extending cytonemes toward the niche (Figure 2B, S1D). We also observed germaria containing cysts with a low pMad level and short, niche-directed cytonemes (Figure S1C), as well as germaria in which GSCs have recolonised the niche (Figure 2C, S1E). These pMad-positive GSCs have cytonemes contacting the niche (Figure 2C, S1E), as observed for non-HS controls (Figure 2A).

In some germaria with cysts, extra somatic cells are found to accumulate in the niche (Figure 2B, S1A-B, G), some of which are pMad-positive (Figure 2B, pink arrowhead) indicating that they are receiving BMP signalling. This is consistent with a previous report of somatic ECs and follicle cell progenitors populating the niche after GSC loss following *hs-bam* expression (36). We hypothesised that accumulation of non-germline cells in the niche could block access for the differentiating cysts preventing them from reoccupying the niche. To test this, we quantitated both the number of niche cells and pMad-positive GCs in germaria undergoing dedifferentiation following a *hs-bam* pulse (Figure S1F-K). These data show that *hs-bam* germaria with pMad-positive GCs have the same number of niche cells as in control non-HS germaria, whereas germaria with extra niche cells contain only pMad-negative GCs (Figure 2D). This dichotomy supports the idea that extra cells in the niche act as a barrier to dedifferentiation.

To study the cytonemes in more detail, we quantitated the number, length and orientation in CBs in the process of niche recolonisation in 3 dpHS *hs-bam* germaria, compared to CBs in control non-HS germaria. These germaria also have germline expression of *LifeAct-tagRFP* and carry a *bam* promoter-GFP (*bamP-GFP*) reporter that marks differentiating cells (Figure 2E, F). The quantitation shows that the dedifferentiating CBs have the same number of cytonemes as the control CBs from non-HS germaria (Figure 2G). However, the cytonemes on dedifferentiating CBs are longer and more niche oriented (Figure 2H-I). To measure cytoneme orientation, a line was drawn from the centre of each GSC toward the niche, with this position marked as 0° on the plots (Figure 2I). This increased length and niche orientation may reflect the increased space available during dedifferentiation when GSCs are lost from the niche. Together, these data show that post-HS, when the niche is empty, the cysts begin to extend cytonemes toward the niche. Over time, if the cysts are not blocked by the accumulation of somatic cells in the niche, they are able to access Dpp, reactivate pMad and dedifferentiate as GSCs (Figure 2J).

The cytonemes on dedifferentiating cells are oriented towards the BMP expressing niche (Figure 2I), consistent with previous reports of cytonemes polarised towards BMP sources in the wing disc (37–39). Therefore, we took a reductionist approach to determine whether a Dpp source is sufficient to induce cytoneme contacts. To this end, we used *Drosophila* S2R+ cells and beads that were either coated with BMP2 or BSA as a negative control. S2 cells have previously been used as a model to study Hedgehog-containing cytonemes (40). First, we tested whether the BMP2 beads were bioactive based on their ability to induce pSmad1/5 activation when added directly to C2C12 cells. These data show that the BMP2 beads activate pSmad1/5, albeit in a lower proportion of cells than observed with soluble BMP2, whereas there is no pSmad1/5 activation in either the control or BSA bead treated cells (Figure S2A-C).

Next, we tested the ability of BMP2 or control BSA beads to induce cytoneme contacts from S2R+ cells that are transfected with expression plasmids encoding the BMP receptor Thickveins (Tkv), as a Tkv-mCherry fusion, and Flag-Mad. Flag-Mad was transfected to increase the cell’s BMP-responsiveness. In the analysis, we included cells that had at least one bead within a distance of 50μm from the centre of their nucleus. These data show that ∼25% of S2R+ cells extend cytoneme contacts to the BMP beads (Figure 2K, M). This is significantly greater than the small proportion of contacts observed with the BSA beads (Figure 2L, M), despite there being no significant difference in the distances between the centre of the cell nucleus and beads across the cells analysed (Figure 2N). Other examples of S2R+ cytoneme contacts with BMP beads and the quantitation from additional biological replicates are shown in Figure S2D-G. Therefore, in this simple system, a bioactive BMP2 bead can induce contacts from Tkv-loaded cytonemes from S2R+ cells. Although this system is heterologous, the findings are broadly relevant to dedifferentiation and other cell fate outcomes by showing that a BMP source can be sufficient for cytonemes to sense and orient towards this source. Together, these *in vitro* and *in vivo* data support a model in which, upon GSC loss, the differentiating cells extend cytonemes to the ovarian niche Dpp, promoting BMP-responsiveness and niche recolonisation.

### Ena is localised at the tips of GSC and CB cytonemes

To test whether cytonemes promote GSC-niche recolonisation by allowing access to niche Dpp, we sought to disrupt GC cytonemes without affecting other critical actin functions. To this end, we explored Ena as a potential cytoneme regulator because it regulates filopodia emergence, without affecting cell division (24, 41). We initially visualised Ena localisation in GCs. Using antibody staining in fixed germaria, we identified Ena in early GCs and at the interface between the CpCs and the GSCs (Figure 3A). Using Vasa staining to mark GCs, Ena was also identified at the tips of GSC cytonemes oriented towards the niche (Figure 3Bi, ii), consistent with its role as an actin polymerase (42, 43). We also detected Ena at the tips of CB cytonemes (Figure 3C) and lamellipodia-like projections (Figure 3Ci, bracket), suggesting that Ena may be a general regulator of GC cytonemes.

**Figure 3:**
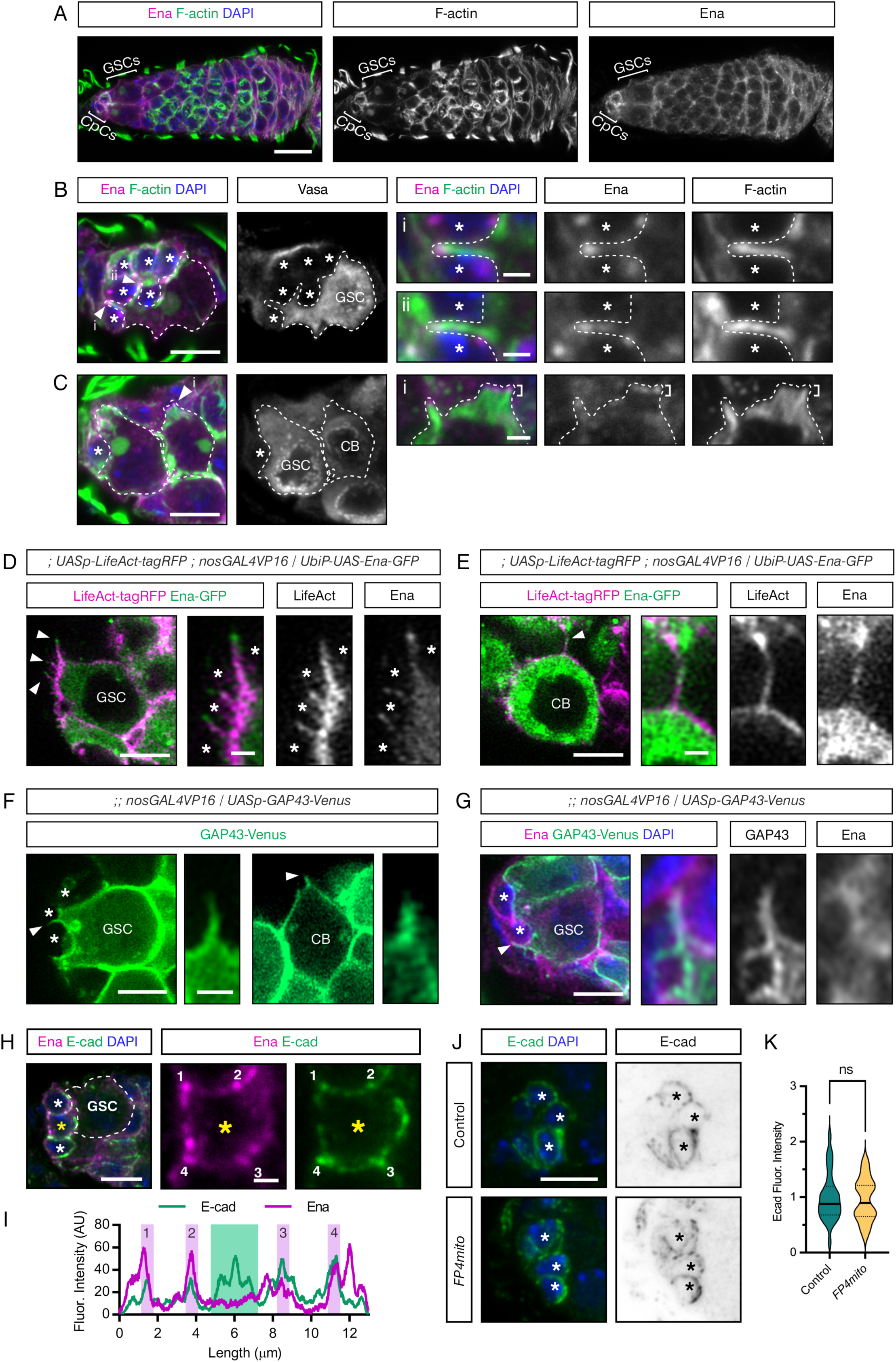
Ena is localised to cytoneme tips. (A) Wildtype germarium immunostained with anti-Ena (magenta), Phalloidin (green) and DAPI (blue). GSCs and CpCs are marked by brackets. Scale bar, 10μm. (B) Immunostaining as in (A) and Vasa (grey inset). A GSC is outlined by a dotted line based on Vasa staining, white arrowheads mark cytonemes (i and ii), that are shown as higher magnification images. (*) Cap Cells. Scale bar, 5μm and (i-ii) 1μm. (C) Immunostaining as in (B). A GSC and CB are outlined by dotted lines. A CB cytoneme is indicated by a white arrowhead and shown magnified in (i). Lamellipodial projections are indicated by brackets. (*) Cap Cells. Scale bar, 5μm and (i) 1μm (D, E) Images of a GSC (D) and CB (E) from live germaria expressing *LifeAct-tagRFP* (magenta and inset) and *Ena-GFP* (green and inset) driven by *nosGAL4VP16.* Higher magnification views of cytonemes are shown. (*) Cap Cells. Scale bar, 5μm and inset 1μm. (F) Image from a live germarium expressing *GAP43-Venus* driven by *nosGAL4VP16.* A GSC and CB are shown, with higher magnification views of the cytonemes. (*) Cap Cells. Scale bar, 5μm and inset 1μm. (G) A GSC imaged in a fixed germarium expressing *GAP43-Venus* driven by *nosGAL4VP16* and immunostained with anti-GFP (green and inset), anti-Ena (magenta and inset) and DAPI (blue). The higher magnification view shows a cytoneme. (*) Cap Cells. Scale bar, 5μm and (inset) 1μm. (H) Wildtype germarium immunostained with anti-Ena (magenta), anti-Ecad (green) and DAPI (blue). The GSC is outlined by a dotted line. (*) Cap Cells. Scale bar, 5μm and inset 1μm. (I) Intensity plot for Ena and Ecad staining shown in (H) across points 1-4 of the CpC marked with a yellow asterisk. (J) Control and *FP4mito* expressing germaria immunostained with anti-Ecad (green and grey inset) and DAPI (blue). (*) Cap Cells. Scale bar, 5μm (K) Intensity plot for Ecad staining, as shown in (J), n=29 and 23, Welch’s unpaired two-tailed t test, p =0.5942 (ns).

To complement these fixed images, we used super-resolution imaging (Zeiss 980, airyscan) to visualise Ena distribution on cytonemes in live *ex vivo* germaria carrying *Ena-GFP* and *LifeAct-tagRFP* reporters activated in the germline by *nosGAL4VP16*. The Ena-GFP transgene encodes a protein fusion driven by the *ubiquitin* promoter with upstream UAS sites. The UAS sites allow stronger expression in the presence of a driver than with the weaker *ubiquitin* promoter alone. Therefore, Ena-GFP expression is specifically enhanced in the germline in the presence of the *nosGal4VP16* driver compared to a control lacking *nosGal4VP16* (Figure S3A). The images from live germaria show that Ena-GFP decorates the length of GSC and CB cytonemes and is concentrated on the tips (Figure 3D, E). Overall, this localisation suggests that Ena may be important for cytoneme extension and, therefore, a potential selective target for manipulating cytoneme behaviour.

As an alternative cytoneme marker to LifeAct, we also used a *UASp-GAP43-Venus* transgene, which revealed protrusions from GSCs extending towards CpCs in a previous study (44). This fusion contains the first 20 amino acids of growth-associated protein 43 (GAP43), which targets Venus to the inner leaflet of the plasma membrane (45). Imaging live germaria expressing *GAP43-Venus* in GCs using *nosGal4VP16* revealed the presence of cytonemes on GSCs and CBs (Figure 3F). In addition, Ena staining in fixed germaria with germline expression of *UASp-GAP43-Venus* reveals that Ena is localised within the cytoneme (Figure 3G). Given that GAP43-Venus is a new marker for studying GC cytonemes, we used the data from live germaria (example in Figure 3F) to quantitate cytoneme properties in GSCs and CBs. These data show that GSCs typically have more cytonemes that are longer and more polarised towards the niche than the CB cytonemes (Figure S3B, C). These findings are consistent with the quantitation for the control CBs (Figure 2G-I) and previous quantitation of GSC and CB cytonemes (19) using *LifeAct-GFP*.

To disrupt Ena function specifically in the ovarian germline, we used an established Ena mislocalisation construct, *FP4mito.* FP4mito contains a multimerised FPPPP (FP4) motif, which is bound by the N-terminal EVH1 domain of Ena with high affinity, fused to a mitochondrial outer membrane targeting signal (27). As targeting the FP4 motif to mitochondria mislocalises Ena there, expression of a *FP4mito* transgene *in vivo* inhibits Ena function (41, 46–48). To specifically mislocalise Ena in GCs, *FP4mito* was expressed in the germline using *nosGAL4VP16* alone or with *LifeAct-GFP* to visualise F-actin. Both the Ena- GFP reporter (Figure S3D) and F-actin (Figure S3E) are observed to accumulate in cytoplasmic aggregates, only in GCs, in *FP4mito* expressing germaria, as described previously for *FP4mito* expressing epidermal cells in the *Drosophila* embryo (41).

Ena can also play a role in cellular adhesion and we observe Ena concentrating at both GSC- CpC and CpC-CpC tricellular junctions (Figure 3H, I), but we do not observe Ena with Ecad at the GSC-niche interface (Figure 3I). Although the data described above show that Ena is mislocalised by germline expression of *FP4mito*, we observe no effect on Ecad levels (Figure 3J, K), suggesting that Ena mislocalisation does not impact GSC-niche adhesion at homeostasis. Overall, these data show that *FP4mito* efficiently mislocalises Ena in the ovarian germline and therefore provides a potential tool for the perturbation of cytoneme formation.

### Ena regulates GC cytoneme formation

Next, we tested the impact of *FP4mito* expression on cytoneme formation by quantifying the number, length and orientation of GSC and CB cytonemes labelled with germline expression of *LifeAct-GFP* in live *ex vivo* germaria (Figure 4A, B). We found that when Ena is mislocalised to mitochondria, there are significantly fewer (∼2-fold) cytonemes on both the GSCs and CBs (Figure 4C). In addition, the mean GSC and CB cytoneme length is significantly reduced by ∼2-fold in *FP4mito* germaria compared with controls (Figure 4D). This is consistent with previous studies where mislocalisation of Ena with *FP4mito* results in a reduced length of filopodia in leading edge cells during *Drosophila* dorsal closure (41).

**Figure 4:**
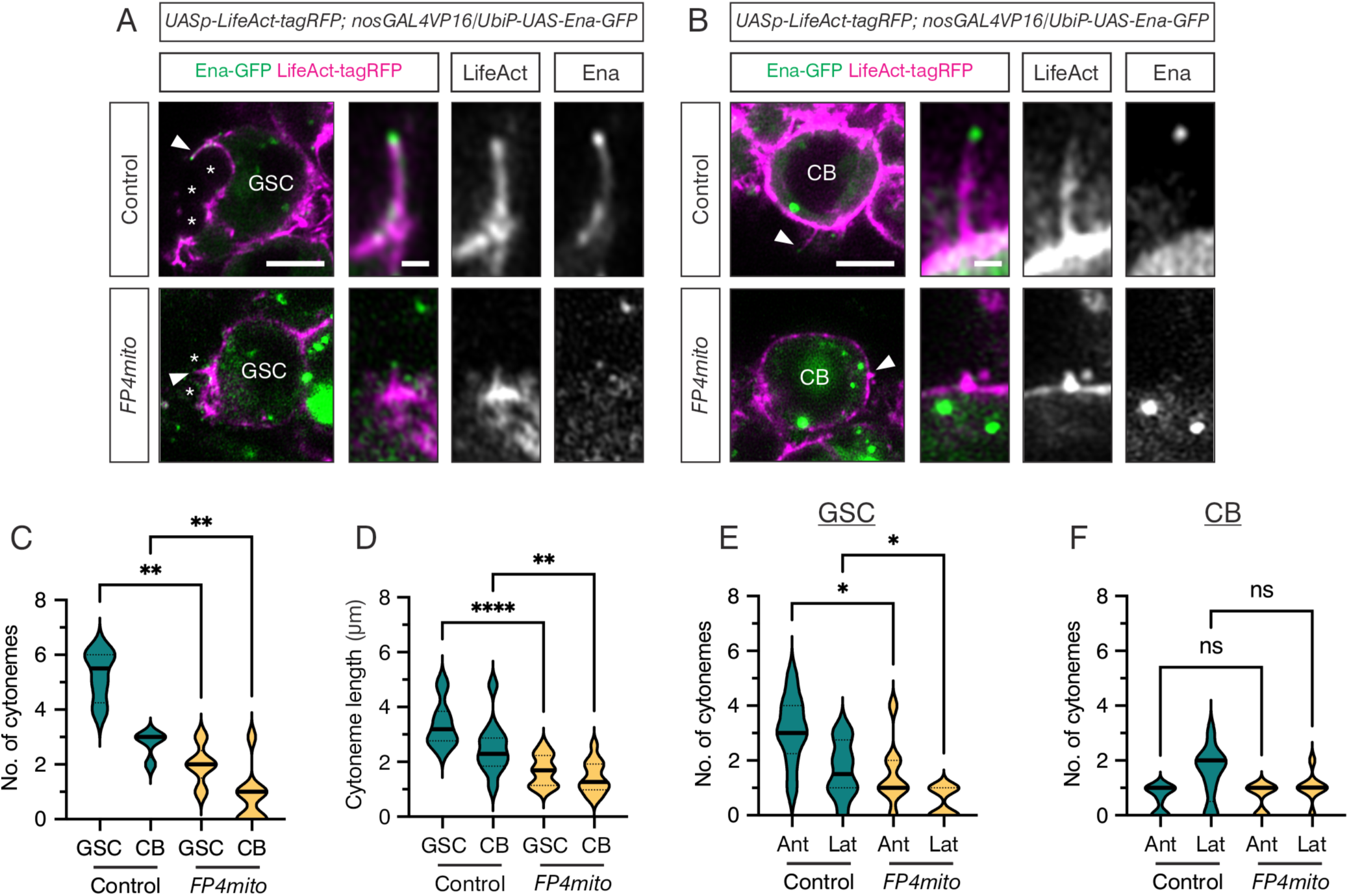
Ena regulates GC cytoneme formation. (A, B) Live images of control and *FP4mito* expressing germaria, with *LifeAct-tagRFP* (magenta and inset) and *Ena-GFP* (green and inset) driven by *nosGAL4VP16*. Images show a GSC (A) or CB (B), with higher magnification views of a cytoneme. (*) Cap Cells. Scale bar, 5μm. (C, D) Graphs showing the number of cytonemes per GSC or CB and cytoneme lengths from live control and *FP4mito* germaria, as shown in (A, B) for representative examples. n = 11-13 cytonemes, Welch’s unpaired t test, **** p = <0.0001 and ** p = <0.01. (E, F) Data from (C, D) showing the number of cytonemes per GC projecting either anteriorly or laterally from live control and *FP4mito* germaria, as shown in (A, B) for representative examples. Welch’s unpaired t test, * p = <0.05.

We next characterised the directionality of GSC cytonemes. Briefly, a line was drawn from the centre of each GSC toward the niche. Cytonemes within a 45° angle from this line were counted as being anteriorly localised, while cytonemes oriented more than 45° were assigned as lateral (19). *FP4mito*-expressing GSCs display a significant shift from anteriorly localised to more lateral cytonemes than controls (Figure 4E), indicating that the directionality of cytoneme formation is perturbed. However, there is no significant change in the orientation of CB cytonemes in *FP4mito* germaria (Figure 4F). In conclusion, Ena mislocalisation perturbs GSC and CB cytoneme formation and those that are formed are shorter and, in the case of GSCs, misoriented relative to the niche.

### Ena mislocalisation impairs BMP signal reception and stem cell fitness

As cytonemes enable GSCs to access niche sequestered Dpp, we tested the effect of perturbing cytoneme formation on BMP signalling and stem cell self-renewal by staining for pMad, Bam, Spectrin and DAPI in *FP4mito* and control germaria. The number of pMad- positive GCs in each background was quantitated. Wildtype germaria typically contain 3-4 pMad-positive GCs, i.e. GSCs and CBs, whereas two or less indicates aberrant differentiation and 5 or more indicates delayed differentiation (19). By 1-week post-eclosion, there is an increase in the number of pMad-positive GCs lost to differentiation in *FP4mito* germaria compared to controls (Figure 5A-C). Furthermore, GSCs expressing *FP4mito* display significantly reduced pMad levels (Figure 5D). Quantitation also reveals that almost all the germaria contain Bam-positive cells or cysts (Figure 5E), showing that the ability of GCs to dedifferentiate is unperturbed by *FP4mito* expression. Together, these data reveal that there is reduced Dpp signal reception and GSC loss in germaria from 1-week old females when cytonemes are disrupted.

**Figure 5:**
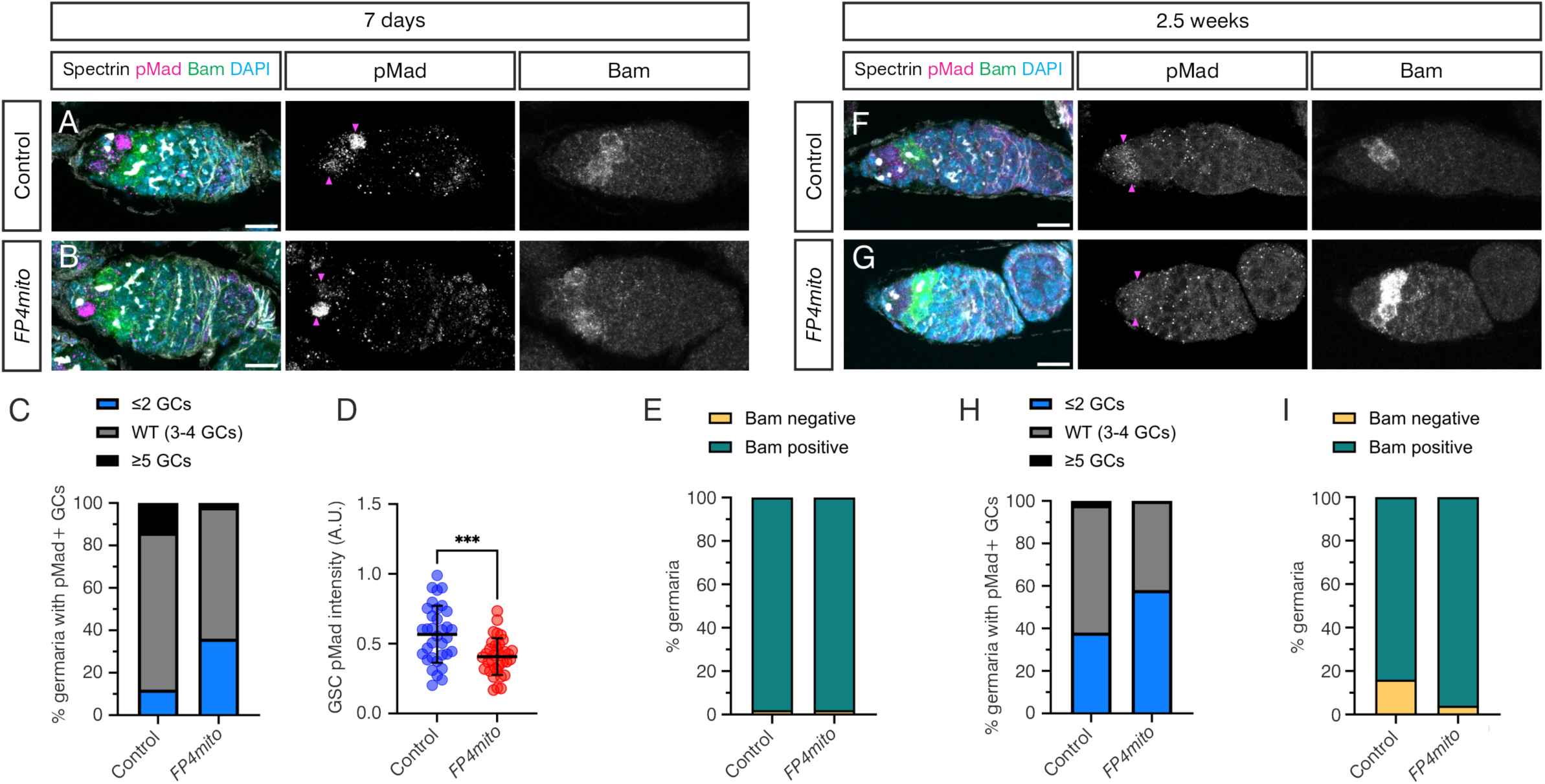
Ena mislocalisation impairs BMP signal reception and stem cell fitness. (A, B) Representative images of control or *FP4mito* germaria aged for 7 days, immunostained with anti-αSpectrin (grey), anti-pMad (magenta and inset), anti-Bam (green and inset) and DAPI (blue). Magenta arrowheads indicate pMad-positive GCs. Scale bar, 10μm. (C) Graph shows the number of pMad-positive GCs counted in 7-day old germaria including the representative germaria shown in (A, B). n = 50. (D) Graph shows the pMad intensity in GSCs in control and *FP4mito* backgrounds. n = 34 per genotype, data shown as mean ± SD, Welch’s two-tailed unpaired t test, p = 0.0003 for comparisons between genotypes. (E) Graph shows the proportion of germaria with and without Bam-positive cysts counted in 7-day old samples. n=50. (F, G) As in (A, B), except the germaria were aged for 2.5 weeks. (H) Graph shows the number of pMad-positive GCs counted in 2.5-week-old germaria, including the representative germaria from (F, G). n = 50. (I) Graph shows the proportion of germaria with and without Bam-positive cysts counted in 2.5-week-old samples. n=50.

By 2.5 weeks post-eclosion, we observe an increase in the percentage of control and *FP4mito* germaria exhibiting aberrant differentiation (Figure 5F-H), relative to 1-week post-eclosion (Figure 5C), consistent with age-related loss. At 2.5 weeks, more *FP4mito* germaria show GSC loss, based on two or less pMad-positive cells (Figure 5H). There is also a small proportion of germaria that are Bam-negative, e.g. that contain 16cc, consistent with age-related decline (Figure 5I). Together, these data show that mislocalisation of Ena perturbs cytoneme formation and the ability of GSCs to respond to niche BMP signalling, resulting in a decline in the stem cell pool.

### Ena mislocalisation impairs GC dedifferentiation

Given that perturbation of cytoneme formation disrupts BMP signal reception, we next used *FP4mito* to test whether cytonemes are necessary for GC dedifferentiation. Specifically, we transiently induced the *hs-bam* transgene as before (Figure 1A), in either a *nosGAL4VP16*>*FP4mito* or *nosGAL4VP16* control background. We also performed a control HS experiment with *nosGAL4VP16*>*FP4mito* or *nosGAL4VP16* flies that lacked the *hs-bam* transgene, using the same HS conditions (Figure S4A-F). Quantitation of these germaria at 1, 3 and 6 dpHS reveals that almost all contain pMad-positive GSCs (Figure S4G), although there is some variation in the relative numbers over time and trends towards GSC loss for *FP4mito* germaria (Figure S4H). Therefore, HS in the absence of a *hs-bam* transgene did not induce ectopic GC differentiation in control or *FP4mito* germaria over the time course (Figure S4I). Consistent with this, most germaria have Bam-positive differentiating cells (Figure S4J), showing that HS in the absence of a *hs-bam* transgene is insufficient to induce aberrant GC differentiation.

Next, we performed the same experiment in control and *FP4mito* germaria that also carried the *hs-bam* transgene. We performed a pair-wise comparison of the number of germaria with pMad-positive GCs (i.e. GSCs or CBs) 1 vs 6 dpHS. On day 1 in both conditions, the majority of GCs are pMad negative and have branched fusomes (Figure 6A, C, E, F). On day 6, a proportion of control *nosGAL4VP16* germaria contain pMad-positive GSCs and a round spectrosome, consistent with niche recolonisation (Figure 6B). Germaria lacking pMad- positive cells and round fusomes were also observed (Figure S5A), indicative of a lack of germline recovery such that all GCs were lost due to differentiation. Quantitation of these data show that ∼40% of control germaria have a significant recovery of pMad-positive GCs (Figure 6E), consistent with the data presented in Figure 1.

**Figure 6:**
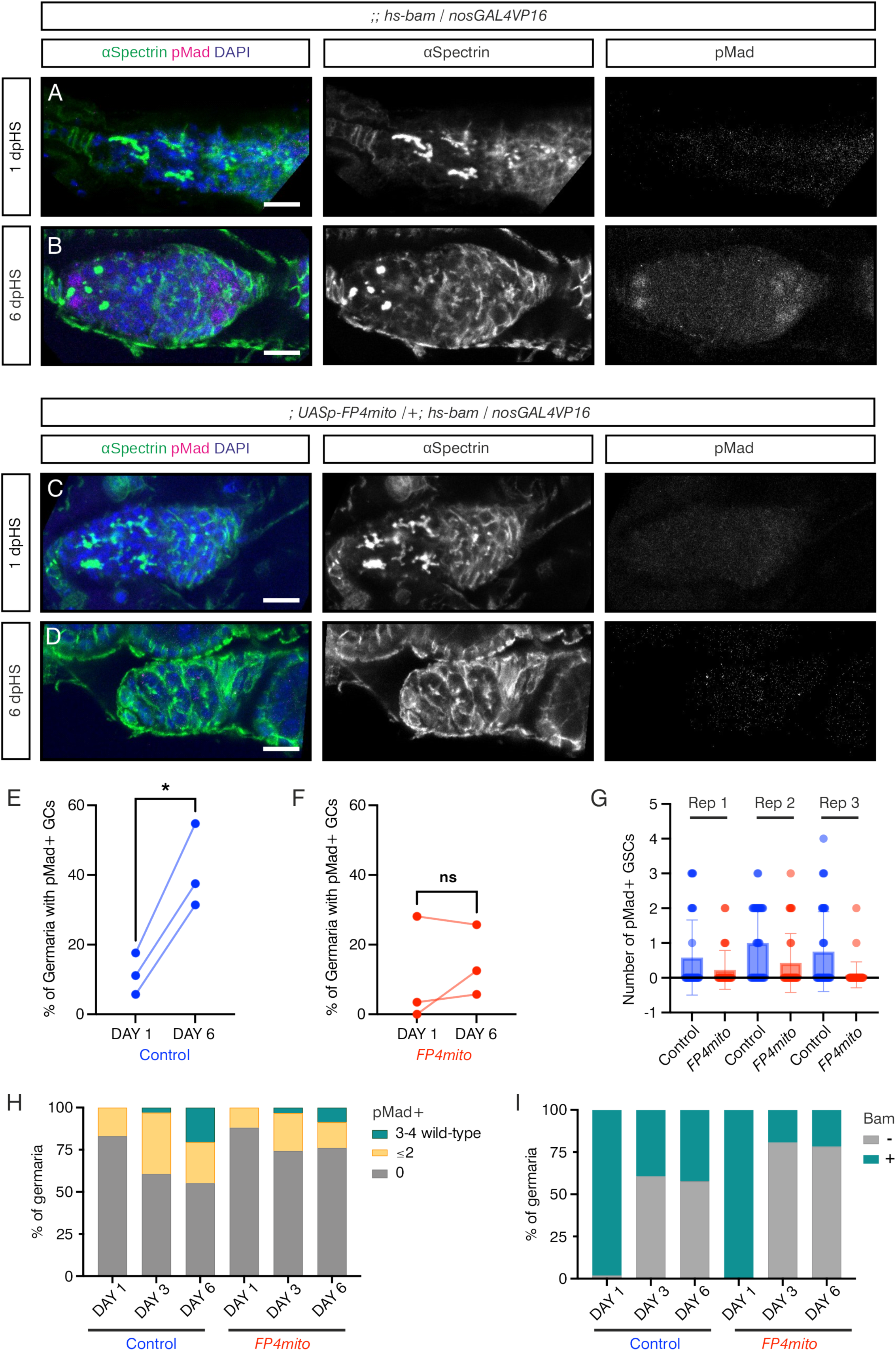
Ena mislocalisation impairs GSC dedifferentiation. (A, B) Control germaria carrying a *hs-bam* transgene and *nosGAL4VP16* immunostained with anti-pMad (magenta and inset), anti-αSpectrin (green and inset) and DAPI (blue). Representative germaria are shown for day 1 (A) and 6 (B) post-HS. Scale bar, 10μm. (C, D) As in (A, B), except germaria also carry a *UASp-FP4mito* transgene. Scale bar, 10μm. (E, F) Quantification of germaria containing pMad-positive early GCs on days 1 and 6 post-HS for (E) control and (F) *FP4mito*. Data are pairwise comparisons, two-tailed paired t test comparing day 1 to day 6, p = 0.0150 for control (*) and p= 0.448 for FP4mito (ns). Each repeat contains at least 30 germaria. (G) Quantification of the number of pMad-positive GSCs per germaria for each repeat. Data shown as mean ± SD. (H, I) Graph shows the number of pMad-positive GCs (H) or Bam-positive GCs (I) counted in 1, 3 and 6 dpHS (representative examples are in Figure S5A-J).

*FP4mito* germaria with either pMad-negative (Figure 6D) or pMad-positive GCs (Figure S5B) were also observed 6 dpHS. However, in contrast to control germaria, there is little or no recovery of pMad-positive GSCs in *FP4mito* expressing germaria (Figure 6F, cf 6E). Additionally control germaria have higher numbers of pMad-positive GSCs compared to *FP4mito* germaria (Figure 6G), indicating that the *FP4mito* GCs are less able to respond to Dpp signalling.

It is possible that less dedifferentiation is observed in *FP4mito* germaria because the cells dedifferentiate normally but are then rapidly lost again to differentiation. To test this, we performed Bam, pMad, Spectrin and DAPI staining on control and *FP4mito* germaria carrying the *hs-bam* transgene, at 1, 3 and 6 dpHS (Figure S5C-L). Quantitation of the number of pMad-positive cells again reveals lower dedifferentiation to pMad-positive cells in the *FP4mito* germaria (Figure 6H, S5M, N). Between days 3 and 6, the proportion of pMad-positive dedifferentiated cells is constant in either the control or *FP4mito* backgrounds, but the cells become more dedifferentiated over time so that there is an increase in the relative proportion of germaria with 3-4 pMad-positive cells (Figure 6H) and GSCs (Figure S5O). These results are inconsistent with dedifferentiation occurring normally in the presence of *FP4mito*, followed by rapid GSC loss, which would result in a reduction in the number of pMad-positive cells at day 6 relative to day 3, rather than an increase.

Similarly, quantitation of Bam-positive cells over the time course reveals that the proportion of differentiating cells is unchanged between days 3 and 6 for each of the control or *FP4mito* backgrounds (Figure 6I). The GCs that do not dedifferentiate are Bam-negative, a characteristic of later stage cysts, consistent with the GCs remaining differentiated. Together, these results indicate that FP4mito, which causes Ena mislocalization and subsequent cytoneme dysfunction, prevents cyst dedifferentiation. These data are consistent with cytonemes playing an important role in the ability of cysts to respond to the self-renewal Dpp signal which is necessary for GCs to recolonise an empty niche.

## Discussion

Here, we provide evidence that actin-rich cytonemes, which are present on all early ovarian GCs (Figures 2E-F, 3B-G, 4) (19), are necessary for robust BMP signal activation and niche occupancy during both homeostasis and dedifferentiation. Our data also show that Ena loss-of-function can be a useful and targeted tool for dysregulating cytoneme function in different contexts. Ena is localized to cytoneme tips, as observed in the filopodia of leading-edge cells during dorsal closure (46). Using a *FP4mito* transgene to mislocalise Ena resulted in less frequent and shorter cytonemes, again consistent with the effects of Ena mislocalisation on filopodia (41, 46). While there are contexts where *FP4mito* expression phenocopies *ena* loss-of-function mutants (41, 48), there are also examples where a stronger or different phenotype is observed with *FP4mito* compared to *ena* mutants (46, 48). This is likely because a subset of cellular Diaphanous (Dia), a processive actin polymerase, can be mislocalised to mitochondria with Ena upon expression of *FP4mito* (46, 47). Indeed, loss of Dia disrupts cytoneme formation in ovarian GSCs and other contexts (5, 19, 39, 49, 50) and both Ena and Dia also have roles in mediating adhesion, with Dia having a role in Ecad endocytosis in epithelial cells (51) and GSCs (19). Bam promotes GC differentiation by the post-transcriptional down-regulation of Ecad (52, 53) which must be re-expressed during dedifferentiation. Therefore, it is possible that disruption to adhesion could impact dedifferentiation by precluding the formation of stable GC-niche interactions. However, *FP4mito* expression does not appear to affect GSC-niche Ecad levels at homeostasis (Fig. 3J-K), suggesting that Dia function is largely unaffected.

We show that cytonemes coordinate niche occupancy and responsiveness to the BMP self-renewal signal during homeostasis and dedifferentiation (Figure 7). While, it would have been interesting to study physiological dedifferentiation, this is a very rare event that occurs at a frequency of only 0.2% (54). Therefore, we relied on inducing a transient *hs-bam* pulse, which is the standard assay in the field for studying dedifferentiation (35, 36, 54). We have also established a tissue culture-based assay for studying BMP cytonemes, as we show that BMP beads can induce contacts by Tkv-loaded cytonemes from S2R+ cells (Figure 2K-N). A similar approach with ESCs and Wnt beads has revealed new insights into cytoneme-Wnt source contacts, including similarities with glutamatergic neuronal synapses (6). An S2 cell model for studying Hh-containing cytonemes revealed that Dispatched stabilises cytonemes and was predictive of their *in vivo* functionality in the *Drosophila* wing disc (40). Therefore, the S2R+ cell cytoneme-BMP bead model will be useful in future studies for dissecting mechanisms and making predictions that can be tested in ovarian GCs and other contexts.

**Figure 7:**
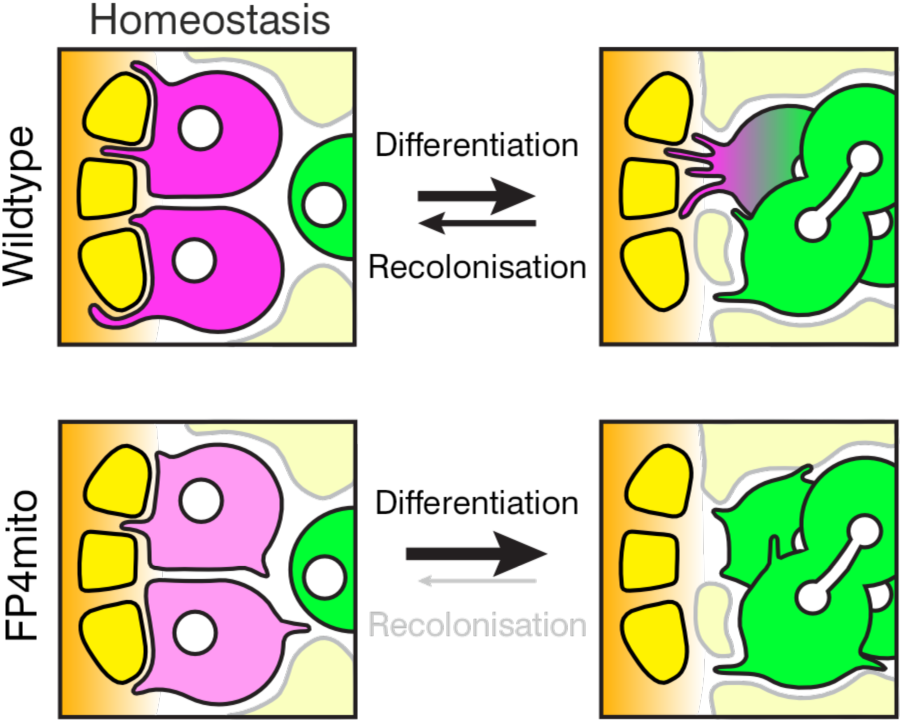
Cytonemes promote stem cell fitness and enable cyst dedifferentation. Model of the role of cytonemes in the ovarian germline at homeostasis, in promoting Dpp signalling and stem cell fitness, and in dedifferentiation, by enabling GCs to access niche-derived Dpp.

Our findings demonstrate a requirement for cytonemes to promote stem cell fitness. While the cytoneme regulators we targeted by knockdown in our previous study affected either cytoneme number or length (19), an advantage of targeting Ena is that both are reduced (Figure 4). Mislocalising Ena reveals that GSCs with disrupted cytonemes have lower BMP signalling and reduced niche occupancy (Figures 5, 7). Cytonemes also direct contact with the BMP source during recolonisation of the niche (Figures 6, 7) and previous studies have shown how cytonemes can orient towards and be stabilised by Dpp-expressing cells (37–39). Moreover, our *in vitro* experiments demonstrate that a BMP source is sufficient to attract and/or stabilise cytonemes, consistent with cytonemes facilitating selective and stable niche interactions. Therefore, our data support cytonemes playing a fundamental role in establishing/maintaining niche interactions and asymmetric signalling, which are central to maintaining the stem cell fate.

Although CBs extend cytonemes in all directions during germline homeostasis, the cytonemes are longer and more oriented towards niche cells during dedifferentiation (Figure 2E-I), which likely reflects in part the space available for cytoneme extension. This cytoneme polarity allows the differentiating cells to become responsive to the self-renewal Dpp signal again, facilitating niche recolonisation and polarised signalling. This is similar to the interplay between cytonemes and stemness during ovarian GSC maintenance, whereby cytonemes allow receipt of the Dpp signal from niche cells, which then promotes the stem cell fate. In further feedback regulation, Dpp signalling induces a second type of GSC cytoneme to establish a Dpp signalling threshold, which allows the daughter that exits the niche to differentiate (19). Similar interdependent relationships between the cause and effect of cytonemes in mediating asymmetric signalling and niche organisation have been described in other contexts. For example, *Drosophila* AMPs extend polarized cytonemes to contact FGF-expressing wing disc niche cells at adherens junctions. This anchors FGF receptor-loaded AMP cytonemes to niche cells, resulting in cytoneme-dependent asymmetric FGF signalling, which reinforces AMP polarity, niche-proximal position and stemness (5). Additionally, *Drosophila* testis GSCs extend cytoneme-like structures called microtubule-dependent nanotubes (MT-nanotubes) into the niche to collect the self-renewal Dpp signal, with Dpp required to induce or stabilise MT-nanotubes (55).

Only a proportion of cysts are capable of dedifferentiating (Figure 1) (54), with 4cc and 8cc previously shown to differentiate (35). As we only detect cytonemes up to the 8cc stage (Figure 2B, S1A), this may explain why dedifferentiation of 16cc does not occur. Therefore, the presence of cytonemes is likely one factor limiting dedifferentiation. Recently, the changes in the transcriptome and translatome during GSC differentiation have been described, and this has revealed two major waves of changes separated by an inflection point around the 4cc stage. From the 8cc stage onwards, gene expression changes associated with terminal differentiation occur (56). This transition is also linked to dramatic changes in actin dynamics. Waves of contractile cortical actin are suppressed in 4-8cc and initiate in 16cc to promote their encapsulation by follicle cells prior to oocyte maturation (57). It is possible that the gene expression and proteome changes that occur in the 8cc prevent the cells from forming cytonemes, further committing the cells to terminal differentiation.

Our data also show that the presence of extra niche cells is associated with a failure to dedifferentiate (Figure 2B, D, S1A, B, G). The additional somatic cells we observed colonising the niche are likely to be ECs, which initially enter the empty niche and respond to Dpp (Figure 2B) (36). These data suggest that the ECs colonising the niche can both form a barrier to GSC-niche contact and outcompete the differentiated cysts for Dpp. Additionally, it is possible that, during homeostasis, niche occupancy by GSCs prevents the CB cytonemes from accessing the niche to ensure that the CBs differentiate rather than dedifferentiate. Based on our findings, we suggest that a general prerequisite for dedifferentiation in diverse contexts is cytoneme-mediated niche contact and that other cells can be a major barrier to achieving this. For example, while *Drosophila* AMPs proximal to the niche extend polarised cytonemes to contact niche cells and receive FGF signals to maintain stemness, more distal AMPs have lateral cytonemes, lose niche contact and differentiate (5). It is possible that, in this context, the presence of proximal AMPs physically blocks distal cells’ cytonemes from contacting the niche, which contributes to these cells remaining in a more differentiated state.

Some ECs express low levels of Dpp, which can weakly promote germline cell dedifferentiation (54). We suggest that soluble Dpp from ECs may initially be able to weakly aid dedifferentiation, but it is insufficient to promote cytoneme contact and BMP responsiveness. In support of this, our data show that cytonemes are oriented towards the niche rather than ECs during dedifferentiation, even when there are extra niche cells present that blocks dedifferentiation (Figure 2B). Similarly, germline cells are largely insensitive to the EC released Dpp (54) and cells that have exited the niche can derepress *bam* transcription and differentiate. Different possibilities exist for the general insensitivity of germline cells to the Dpp secreted by ECs. Firstly, the Dpp level may be too low. Secondly, the insensitivity may reflect that ECs lack the heparan sulphate proteoglycan Dally (58, 59), which binds Dpp (60). Previously, we showed that Tkv activation on cytonemes occurs at sites where Dally-Dpp is concentrated, suggesting that Dally may present Dpp to the cytoneme (19). It is possible that in the absence of Dally presentation, cytonemes cannot recognise the soluble Dpp released by ECs. This would be similar to the role of the Glycosylphosphatidylinositol (GPI) anchor in presenting the FGF ligand to air sac primordium (ASP) cytonemes for contact-dependent release and signal reception. In contrast, the ASP cytonemes cannot respond to secreted FGF ligand (61). Thirdly, it is possible that the germline cells respond to the EC Dpp in a cytoneme- independent manner resulting in a qualitatively different response to cytoneme-mediated Dpp reception, which cannot support dedifferentiation. Potential support for this comes from data from the male germline showing that soluble Dpp elicits a different signalling response to the contact-dependent response to niche Dpp (62).

Stem cell maintenance and dedifferentiation are critical for tissue damage repair and underpin the regenerative medicine strategies that rely on stem cell treatments or cellular reprogramming to induced pluripotent stem cells (63–65). As such, manipulating cytonemes could improve regenerative medicine outcomes by favouring particular cell fate outcomes. In addition, defective stem cell maintenance and dedifferentiation is associated with diseases, including ageing-related diseases (66), neurodevelopmental disorders (64, 67) and different cancer types (68, 69). Therefore, cytonemes may represent a new potential therapeutic target for treating the broad spectrum of diseases caused by defective stem cell maintenance and dedifferentiation.

## Materials and Methods

### Fly Stocks

All crosses were undertaken at 25°C on standard fly media (yeast 50g/L, glucose 78g/L, maize 72g/L, agar 8g/L, 10% (v/v) nipagin in ethanol 27ml/L and propionic acid 3ml/L). Stocks used are listed in Table S1. Transgenic line generation is described in the SI Methods.

### Heat shock

Flies 1-3 days post eclosion were heat shocked in vials containing Whatman paper to absorb condensation. Vials were heat shocked in a water bath for 1.5 hours at 37°C and then returned to 25°C for 1 hour before a second 1.5 hours heat shock at 37°C. Flies were then transferred to a new vial and returned to 25°C for the duration of the experiment.

### Immunofluorescence

For standard antibody stain, ovaries were dissected in PBS and fixed in 4% formaldehyde for 20 minutes at room temperature. Ovaries were blocked in 5x Western Blocking buffer (Merck) or 10% BSA in PBST (0.1% Tergitol 15-S-9) for 1 hour and then incubated with primary antibodies (Table S3) at 4°C overnight. They were then washed in PBSTw (0.1% Tween 20 in PBS) before incubation at room temperature for 2 hours in 0.1x WB with secondary antibodies. Phalloidin (0.033μM) and DAPI staining was carried out in the penultimate wash for 30 minutes at room temperature and ovaries were mounted in Prolong Diamond or Gold (Invitrogen).

For staining of cytonemes, ovaries were dissected in PEM buffer (80mM PIPES, 5mM EDTA, 1mM Magnesium chloride, pH7.4 adjusted with NaOH) and fixed immediately in PEM with 4% formaldehyde, 5μM paclitaxel (Merck), 1μM calcium chloride for 30 minutes at room temperature. Washes were carried out in PBST (0.1% Tergitol 15-S-9) and ovaries were then processed as above.

To use anti-Bam and anti-αSpectrin together, anti-αSpectrin was labelled with Zenon AlexaFluor 555 (Thermo Fisher Scientific, according to the manufacturer’s instructions) and added post-secondary antibody incubation and washes. Ovaries were incubated with Zenon labelled anti-αSpectrin for 1 hour at room temperature.

### Live imaging

Live imaging was performed as described (70).

Imaging settings, quantification and statistical tests are described in the SI Methods.

### S2R+ Cell Culture and Immunofluorescence

S2R+ cells (DGRC, RRID:CVCL_Z831) were maintained at 25°C in Schneider’s Drosophila Medium (Gibco) supplemented with 10% foetal bovine serum (FBS; Gibco) and 1% penicillin-streptomycin (Merck) and plated onto poly-L-lysine (Merck) coated coverslips at 0.75 x 10^6^ density. Cells (0.75 x 10^6^) were co-transfected with 200ng pAc-Tkv-mCherry (Thickveins C-terminal mCherry fusion) and pAc-Mad-FLAG (71) plasmids each using Effectene reagent (Qiagen). After a 72-hour incubation, cells were resuspended in serum-free media and plated onto poly-L-lysine (Merck) coated coverslips at a 0.75 x 10^6^ density. They were serum-starved for 1 hour before addition of serum-free medium containing 3 x 10^5^ BSA- or BMP2-conjugated beads (see below). The cells were incubated for 3 hours at 25°C before fixation (40). Samples were permeabilised for 1 hour at room temperature in 0.1% Igepal CA-630 (Merck), 5% BSA (Merck), 0.3M glycine (Fisher Scientific, CAS: 56-40-6) in 1xPBS, followed by overnight incubation at 4°C of primary antibodies in 1% BSA in PBSTw (0.1% Tween 20 in PBS) (Table S3). Cells were washed four times in PBS and incubated for 2 hours with secondary antibodies in 1% BSA in PBSTw. Samples were washed twice in PBSTw for 10 minutes and DAPI staining was carried out in the penultimate wash. Coverslips were further washed twice in PBSTw before mounting in ProLong Diamond (Invitrogen). Only cells with Tkv-mCherry fluorescence and beads within 50µm of the nucleus were imaged. The bead to nucleus distance was calculated for all beads close to a cell if the distance was within 50µm.

### Bead conjugation

BSA and Human/Mouse/Rat BMP-2 Recombinant Protein (Gibco, 120-02C-10UG) were immobilised onto 2.8 µm Dynabeads M-270 carboxylic acid (Invitrogen) as previously described (72). Briefly, 30µg beads were activated by a 30-minute incubation with carbodiimide (Merck) and N-hydroxyl succinamide (Merck). Following activation, beads were washed three times with 25 mM cold 2-(N-morpholino) ethanesulfonic acid (MES; Merck) buffer (pH5), then incubated for 1 hour with either 0.1% BSA in PBSTw (0.1% Tween 20 in PBS) or 500ng BMP2 (Gibco, 120-02C-10UG) in 25mM MES buffer. The beads were washed with PBS before resuspending them in 1% BSA PBS and stored at 4°C.

### C2C12 Cell Culture, Immunofluorescence and pSmad1/5 quantitation

C2C12 cells were maintained in DMEM high-glucose media (Sigma) supplemented with 2 mM L-glutamine (Gibco), and 10% FBS (Gibco) at 37 °C. Cells were seeded at a density of 1 x 10^5^ cells/well on coverslips. After 6 hours, the cells were serum-starved overnight. The cells were then incubated in serum-free DMEM medium (Sigma) with 11nM soluble BMP2 (Gibco, 120- 02C-10UG) or 3 x 10^5^ BSA- or BMP2-conjugated beads. Cells were incubated at 37°C for 2 hours before being washed in PBS and fixed with 4% PFA for 20 minutes. Samples were permeabilised in 0.2 M glycine and 0.5% Triton X-100 in PBS for 30 minutes and blocked using 2% fish skin gelatine in PBS. Cells were incubated at room temperature with anti- pSmad1/5 (Table S3) for 1 hour, washed in PBS, and incubated with secondary antibody for 30 minutes. Antibodies were diluted in 2% fish skin gelatine in PBS. Samples were washed in PBS and mounted in ProLong Gold with DAPI (ThermoFisher Scientific). pSmad1/5 quantitation was performed as described (73), with an appropriate intensity threshold for each replicate. For the bead experiments, only cells in contact with a bead were included in the quantitation.

All study data are included in the article and/or supporting information.

## Acknowledgements

We thank Tom Millard for suggesting Ena as a target, and Sanjai Patel, the University of Manchester Fly Facility and Bioimaging Facility for support. This work was supported by a Biotechnology and Biological Sciences Research Council project grant (BB/V015060/1) to H.L.A.

## Notes

### Competing Interest Statement

The authors have declared no competing interest.

### Summary of Updates

This version has been revised to include minor text changes.

## References

1. P. Rojas-Rios, A. Gonzalez-Reyes, Concise review: The plasticity of stem cell niches: a general property behind tissue homeostasis and repair. Stem Cells 32, 852–859 (2014).

2. F. A. Ramirez-Weber, T. B. Kornberg, Cytonemes: cellular processes that project to the principal signaling center in Drosophila imaginal discs. Cell 97, 599–607 (1999).

3. C. A. Daly, E. T. Hall, S. K. Ogden, Regulatory mechanisms of cytoneme-based morphogen transport. Cell Mol Life Sci 79, 119 (2022).

4. P. Ramesh, N. S. Dey, A. Kanwal, S. Mandal, L. Mandal, Relish plays a dynamic role in the niche to modulate Drosophila blood progenitor homeostasis in development and infection. Elife 10 (2021).

5. A. Patel et al., Cytonemes coordinate asymmetric signaling and organization in the Drosophila muscle progenitor niche. Nat Commun 13, 1185 (2022).

6. S. Junyent et al., Specialized cytonemes induce self-organization of stem cells. Proc Natl Acad Sci U S A 117, 7236–7244 (2020).

7. S. Junyent, J. Reeves, E. Gentleman, S. J. Habib, Pluripotency state regulates cytoneme selectivity and self-organization of embryonic stem cells. J Cell Biol 220 (2021).

8. T. Xie, A. C. Spradling, A niche maintaining germ line stem cells in the Drosophila ovary. Science 290, 328–330 (2000).

9. D. Kirilly, T. Xie, The Drosophila ovary: an active stem cell community. Cell Res 17, 15–25 (2007).

10. T. Xie, A. C. Spradling, decapentaplegic is essential for the maintenance and division of germline stem cells in the Drosophila ovary. Cell 94, 251–260 (1998).

11. X. Song et al., Bmp signals from niche cells directly repress transcription of a differentiation-promoting gene, bag of marbles, in germline stem cells in the Drosophila ovary. Development 131, 1353–1364 (2004).

12. X. Song, C. H. Zhu, C. Doan, T. Xie, Germline stem cells anchored by adherens junctions in the Drosophila ovary niches. Science 296, 1855–1857 (2002).

13. D. Chen, D. McKearin, Dpp signaling silences bam transcription directly to establish asymmetric divisions of germline stem cells. Curr Biol 13, 1786–1791 (2003).

14. D. M. McKearin, A. C. Spradling, bag-of-marbles: a Drosophila gene required to initiate both male and female gametogenesis. Genes Dev 4, 2242–2251 (1990).

15. D. McKearin, B. Ohlstein, A role for the Drosophila bag-of-marbles protein in the differentiation of cystoblasts from germline stem cells. Development 121, 2937–2947 (1995).

16. H. Lin, L. Yue, A. C. Spradling, The Drosophila fusome, a germline-specific organelle, contains membrane skeletal proteins and functions in cyst formation. Development 120, 947–956 (1994).

17. M. de Cuevas, A. C. Spradling, Morphogenesis of the Drosophila fusome and its implications for oocyte specification. Development 125, 2781–2789 (1998).

18. S. G. Wilcockson, C. Sutcliffe, H. L. Ashe, Control of signaling molecule range during developmental patterning. Cell Mol Life Sci 74, 1937–1956 (2017).

19. S. G. Wilcockson, H. L. Ashe, Drosophila Ovarian Germline Stem Cell Cytocensor Projections Dynamically Receive and Attenuate BMP Signaling. Dev Cell 50, 296–312 e295 (2019).

20. P. Rojas-Rios, I. Guerrero, A. Gonzalez-Reyes, Cytoneme-mediated delivery of hedgehog regulates the expression of bone morphogenetic proteins to maintain germline stem cells in Drosophila. PLoS Biol 10, e1001298 (2012).

21. M. R. Mejillano et al., Lamellipodial versus filopodial mode of the actin nanomachinery: pivotal role of the filament barbed end. Cell 118, 363–373 (2004).

22. C. Lebrand et al., Critical role of Ena/VASP proteins for filopodia formation in neurons and in function downstream of netrin-1. Neuron 42, 37–49 (2004).

23. M. Barzik et al., Ena/VASP proteins enhance actin polymerization in the presence of barbed end capping proteins. J Biol Chem 280, 28653–28662 (2005).

24. J. E. Bear et al., Antagonism between Ena/VASP proteins and actin filament capping regulates fibroblast motility. Cell 109, 509–521 (2002).

25. K. Rottner, B. Behrendt, J. V. Small, J. Wehland, VASP dynamics during lamellipodia protrusion. Nat Cell Biol 1, 321–322 (1999).

26. T. M. Svitkina et al., Mechanism of filopodia initiation by reorganization of a dendritic network. J Cell Biol 160, 409–421 (2003).

27. J. E. Bear et al., Negative regulation of fibroblast motility by Ena/VASP proteins. Cell 101, 717–728 (2000).

28. L. Pasic, T. Kotova, D. A. Schafer, Ena/VASP proteins capture actin filament barbed ends. J Biol Chem 283, 9814–9819 (2008).

29. F. Kang, R. O. Laine, M. R. Bubb, F. S. Southwick, D. L. Purich, Profilin interacts with the Gly-Pro-Pro-Pro-Pro-Pro sequences of vasodilator-stimulated phosphoprotein (VASP): implications for actin-based Listeria motility. Biochemistry 36, 8384–8392 (1997).

30. A. Schirenbeck et al., The bundling activity of vasodilator-stimulated phosphoprotein is required for filopodium formation. Proc Natl Acad Sci U S A 103, 7694–7699 (2006).

31. D. A. Applewhite et al., Ena/VASP proteins have an anti-capping independent function in filopodia formation. Mol Biol Cell 18, 2579–2591 (2007).

32. J. A. Scott et al., Ena/VASP proteins can regulate distinct modes of actin organization at cadherin-adhesive contacts. Mol Biol Cell 17, 1085–1095 (2006).

33. H. Sano et al., The Drosophila actin regulator ENABLED regulates cell shape and orientation during gonad morphogenesis. PLoS One 7, e52649 (2012).

34. B. Ohlstein, D. McKearin, Ectopic expression of the Drosophila Bam protein eliminates oogenic germline stem cells. Development 124, 3651–3662 (1997).

35. T. Kai, A. Spradling, Differentiating germ cells can revert into functional stem cells in Drosophila melanogaster ovaries. Nature 428, 564–569 (2004).

36. T. Kai, A. Spradling, An empty Drosophila stem cell niche reactivates the proliferation of ectopic cells. Proc Natl Acad Sci U S A 100, 4633–4638 (2003).

37. F. Hsiung, F. A. Ramirez-Weber, D. D. Iwaki, T. B. Kornberg, Dependence of Drosophila wing imaginal disc cytonemes on Decapentaplegic. Nature 437, 560–563 (2005).

38. S. Roy, F. Hsiung, T. B. Kornberg, Specificity of Drosophila cytonemes for distinct signaling pathways. Science 332, 354–358 (2011).

39. S. Roy, H. Huang, S. Liu, T. B. Kornberg, Cytoneme-mediated contact-dependent transport of the Drosophila decapentaplegic signaling protein. Science 343, 1244624 (2014).

40. W. J. Bodeen, S. Marada, A. Truong, S. K. Ogden, A fixation method to preserve cultured cell cytonemes facilitates mechanistic interrogation of morphogen transport. Development 144, 3612–3624 (2017).

41. J. Gates et al., Enabled plays key roles in embryonic epithelial morphogenesis in Drosophila. Development 134, 2027–2039 (2007).

42. J. D. Winkelman, C. G. Bilancia, M. Peifer, D. R. Kovar, Ena/VASP Enabled is a highly processive actin polymerase tailored to self-assemble parallel-bundled F-actin networks with Fascin. Proc Natl Acad Sci U S A 111, 4121–4126 (2014).

43. D. Breitsprecher et al., Clustering of VASP actively drives processive, WH2 domain-mediated actin filament elongation. EMBO J 27, 2943–2954 (2008).

44. T. Panchal et al., Specification and spatial arrangement of cells in the germline stem cell niche of the Drosophila ovary depend on the Maf transcription factor Traffic jam. PLoS Genet 13, e1006790 (2017).

45. M. Mavrakis, R. Rikhy, J. Lippincott-Schwartz, Plasma membrane polarity and compartmentalization are established before cellularization in the fly embryo. Dev Cell 16, 93–104 (2009).

46. C. C. Homem, M. Peifer, Exploring the roles of diaphanous and enabled activity in shaping the balance between filopodia and lamellipodia. Mol Biol Cell 20, 5138–5155 (2009).

47. S. H. Nowotarski, N. McKeon, R. J. Moser, M. Peifer, The actin regulators Enabled and Diaphanous direct distinct protrusive behaviors in different tissues during Drosophila development. Mol Biol Cell 25, 3147–3165 (2014).

48. J. Gates et al., Enabled and Capping protein play important roles in shaping cell behavior during Drosophila oogenesis. Dev Biol 333, 90–107 (2009).

49. S. Fereres, R. Hatori, M. Hatori, T. B. Kornberg, Cytoneme-mediated signaling essential for tumorigenesis. PLoS Genet 15, e1008415 (2019).

50. H. Huang, T. B. Kornberg, Myoblast cytonemes mediate Wg signaling from the wing imaginal disc and Delta-Notch signaling to the air sac primordium. Elife 4, e06114 (2015).

51. R. Levayer, A. Pelissier-Monier, T. Lecuit, Spatial regulation of Dia and Myosin-II by RhoGEF2 controls initiation of E-cadherin endocytosis during epithelial morphogenesis. Nat Cell Biol 13, 529–540 (2011).

52. R. Shen, C. Weng, J. Yu, T. Xie, eIF4A controls germline stem cell self-renewal by directly inhibiting BAM function in the Drosophila ovary. Proc Natl Acad Sci U S A 106, 11623–11628 (2009).

53. Z. Jin et al., Differentiation-defective stem cells outcompete normal stem cells for niche occupancy in the Drosophila ovary. Cell Stem Cell 2, 39–49 (2008).

54. Z. Liu et al., Coordinated niche-associated signals promote germline homeostasis in the Drosophila ovary. J Cell Biol 211, 469–484 (2015).

55. M. Inaba, M. Buszczak, Y. M. Yamashita, Nanotubes mediate niche-stem-cell signalling in the Drosophila testis. Nature 523, 329–332 (2015).

56. T. J. Samuels, J. Gui, D. Gebert, F. Karam Teixeira, Two distinct waves of transcriptome and translatome changes drive Drosophila germline stem cell differentiation. EMBO J 43, 1591–1617 (2024).

57. S. Chanet, J. R. Huynh, Collective Cell Sorting Requires Contractile Cortical Waves in Germline Cells. Curr Biol 30, 4213–4226 e4214 (2020).

58. C. Y. Tseng et al., Smad-Independent BMP Signaling in Somatic Cells Limits the Size of the Germline Stem Cell Pool. Stem Cell Reports 11, 811–827 (2018).

59. M. Liu, T. M. Lim, Y. Cai, The Drosophila female germline stem cell lineage acts to spatially restrict DPP function within the niche. Sci Signal 3, ra57 (2010).

60. T. Akiyama et al., Dally regulates Dpp morphogen gradient formation by stabilizing Dpp on the cell surface. Dev Biol 313, 408–419 (2008).

61. L. Du, A. Sohr, Y. Li, S. Roy, GPI-anchored FGF directs cytoneme-mediated bidirectional contacts to regulate its tissue-specific dispersion. Nat Commun 13, 3482 (2022).

62. S. M. Ridwan et al., Diffusible fraction of niche BMP ligand safeguards stem-cell differentiation. Nat Commun 15, 1166 (2024).

63. C. Jopling, S. Boue, J. C. Izpisua Belmonte, Dedifferentiation, transdifferentiation and reprogramming: three routes to regeneration. Nat Rev Mol Cell Biol 12, 79–89 (2011).

64. A. Brunet, M. A. Goodell, T. A. Rando, Ageing and rejuvenation of tissue stem cells and their niches. Nat Rev Mol Cell Biol 24, 45–62 (2023).

65. K. Takahashi, S. Yamanaka, A decade of transcription factor-mediated reprogramming to pluripotency. Nat Rev Mol Cell Biol 17, 183–193 (2016).

66. L. B. Boyette, R. S. Tuan, Adult Stem Cells and Diseases of Aging. J Clin Med 3, 88–134 (2014).

67. A. B. Caldwell et al., Dedifferentiation and neuronal repression define familial Alzheimer’s disease. Sci Adv 6 (2020).

68. X. Chu et al., Cancer stem cells: advances in knowledge and implications for cancer therapy. Signal Transduct Target Ther 9, 170 (2024).

69. S. Schwitalla et al., Intestinal tumorigenesis initiated by dedifferentiation and acquisition of stem-cell-like properties. Cell 152, 25–38 (2013).

70. S. G. Wilcockson, H. L. Ashe, Live imaging of the Drosophila ovarian germline stem cell niche. STAR Protoc 2, 100371 (2021).

71. B. Muller, B. Hartmann, G. Pyrowolakis, M. Affolter, K. Basler, Conversion of an extracellular Dpp/BMP morphogen gradient into an inverse transcriptional gradient. Cell 113, 221–233 (2003).

72. M. Lowndes, S. Junyent, S. J. Habib, Constructing cellular niche properties by localized presentation of Wnt proteins on synthetic surfaces. Nat Protoc 12, 1498–1512 (2017).

73. T. Malinauskas et al., Molecular mechanism of BMP signal control by Twisted gastrulation. Nat Commun 15, 4976 (2024).

