## Supplementary Information for "Ovarian germline stem cell dedifferentiation is cytoneme dependent"

**This PDF file includes:**

Materials and Methods
Figures S1 to S5
Tables S1 to S3
SI References

**Materials and Methods**

**Transgenic line generation**

To generate *pUASp-FP4mito-Halo7-attB, FP4mito* was amplified from genomic DNA from the line *y[1] w[*]; P{w[+mC]=UAS-FLAG-HA-FP4mito}3* (Bloomington Cat#58481; RRID:BDSC_58481) and *Halo7* was amplified (Table S2) from *UAS-7xHalo7::CAAX* (Addgene #87647) (1) and inserted into *pUASP-attB* (DGRC Stock 1358; RRID:DGRC_1358) digested with XbaI and KpnI. *pUASp-FP4mito-Halo7-attB* was microinjected into *y w M(eGFP, vas-int, dmRFP)ZH-2A; P{CaryP}attP40* (Stock 13-20; University of Cambridge Fly Facility) using *phiC31* mediated transgenesis at the University of Manchester Fly Facility.

**Imaging Settings**

Imaging was performed on a Leica SP8 inverted microscope with either a HC PL APO CS2 40x/1.30 oil immersion objective or HC PL APO CS2 motCORR (Water) immersion objective (as in (2)) or a Zeiss 980 Axio Observer 7 with a Plan-Apochromat 63x/1.40 Oil DIC M27 objective.

**Confocal imaging on LSM 980**

Images were collected on a Zeiss LSM 980 Axio Observer 7 with a Plan-Apochromat 63x/1.40 Oil DIC M27 objective The confocal settings were as follows, pinhole (0.76AU/0.86AU/1AU), LSM scan speed 8, bidirectional, 2x line averaging and image size 1884x1884. Z-step sizes were system optimised. Images were collected using GaAsP-PMT and Multialkali-PMT detectors with the following detection settings; DAPI 408-502nm, AF488 (508-544nm)*;* AF555 (552-632nm)*;* and AF647 (642-757nm) with mirrors MBS 488/561/639 and MBS405 and using the 405 (0.2%) 488nm (2%), 561nm (0.5%) and 639nm (4%) laser lines respectively. Images were collected sequentially where it was not possible to eliminate cross-over between channels. Images were processed with LSM Plus Processing (Zeiss). For whole germarium images maximum projections of 5-10 slices are shown in the figures.

Additional modifications included pinhole (0.84AU/1AU), LSM scan speed 12, 5x zoom, 4x line averaging and image size 1512x1512. With additional detector Extended Red GaAsP-PMT and alternate laser powers: 488nm (0.2%), 639nm (4%) 402nm (0.2%) 561nm (0.2%).

For imaging with the near-infrared channel included the following settings were used: Images were collected on a Zeiss LSM 980 Axio Observer 7 with a Plan-Apochromat 63x/1.40 Oil DIC M27 objective The confocal settings were as follows, pinhole (0.88AU/1AU/1.16AU), LSM scan speed 14, 4x zoom, bidirectional, image size 1116x1116 and 8x line averaging. Z-step sizes were system optimised. Images were collected using GaAsP-PMT, Extended Red GaAsP-PMT and Multialkali-PMT detectors with the following detection settings; DAPI (408-

502nm); AF488 (508-544nm); AF555 (552-632nm); AF647 (642-736nm) and AF790 (743-870nm) with mirrors MBS 488/561/639 and MBS405/730 and using the 405 (0.5%) 488nm (1.5%), 561nm (0.2%), 639nm (4%) and 730nm (2%) laser lines respectively. Images were collected sequentially where it was not possible to eliminate cross-over between channels.

**Super Resolution Imaging**

Images were collected on a Zeiss LSM 980 Axio Observer 7 with a LD LCI Plan-Apochromat 40x/1.2 Imm Korr DIC M27 FCS objective with 3 to 6x zoom. The settings were as follows, pinhole 5AU, LSM scan speed 8 or 9, bidirectional with Airyscan Super Resolution mode. Number of Z sections were system optimised. Format and laser and detector settings were system optimised for Super Resolution. Images were collected using an Airyscan 2 detector GaAsP-PMT with the following spectral channel settings: eGFP (420-480nm, 495-550nm), Venus (380-735nm) or tagRFP (380-735nm) using the 488nm (1 or 2%) or 561nm (0.5%) laser lines respectively. Images were processed using LSM Airyscan processing (Zeiss). Maximum intensity projections and single slices of these stacks are shown in the figures.

**Quantification**

Cytoneme length, number and orientation were measured on Image J as described (2).
pMad intensity was measured by taking the Integrated Density (IntDen) of the pMad-positive cell on Image J using the measure plugin. It was background corrected and normalised to the DAPI stain within the same area, as indicated below. 32 GSCs were analysed per genotype.

Normalised pMad Intensity = pMad IntDen – (Area x Mean background)/ DAPI IntDen

Ecad intensity was measured on Image J using the measure plugin and plotted on GraphPad Prism (RRID:SCR_002798).

**Statistical Tests**

Images were processed using Fiji (3). Graphs were plotted and analysis was performed on GraphPad Prism (RRID:SCR_002798).

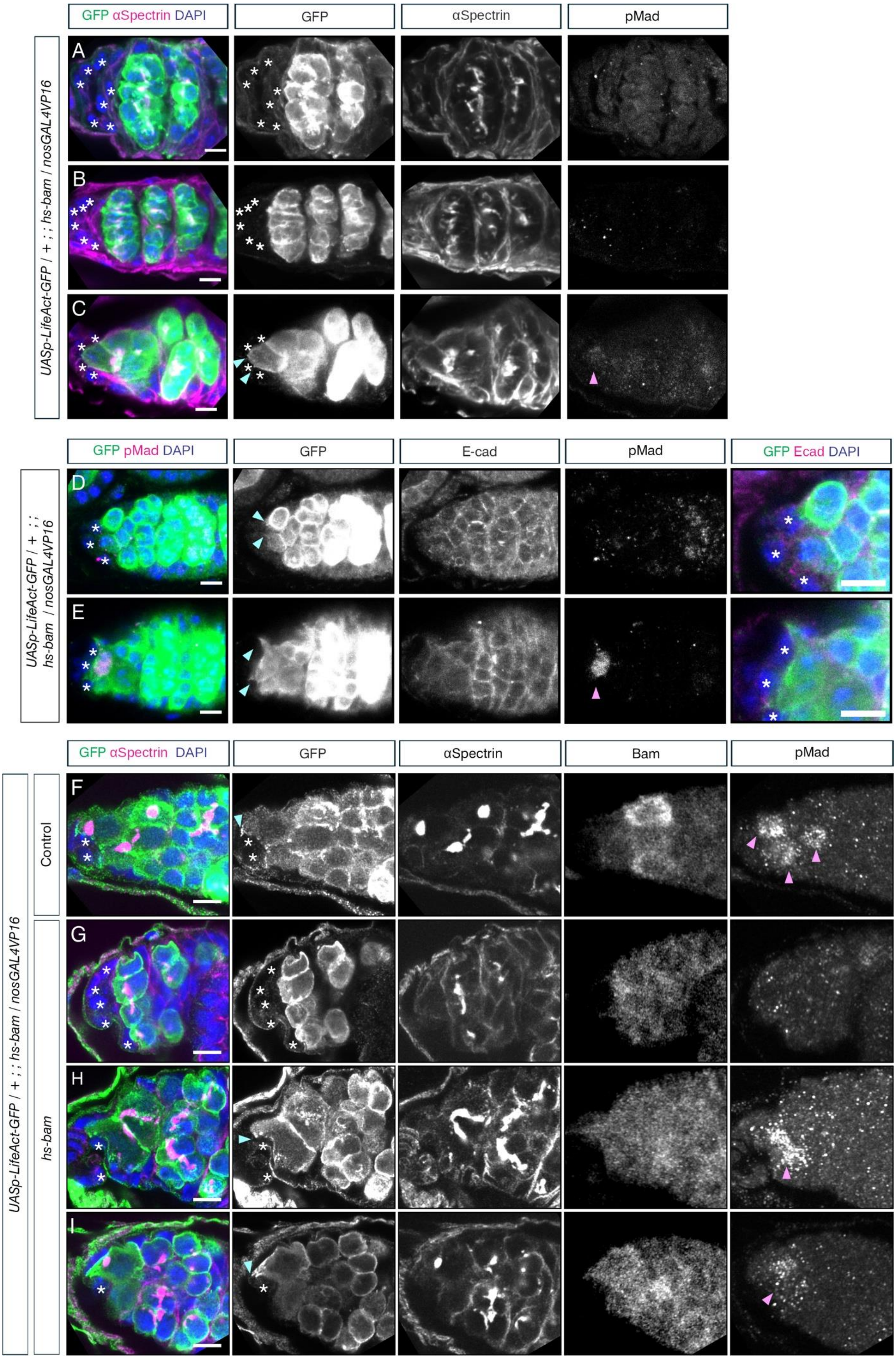

GFP αSpectrin DAPI
GFP
αSpectrin
pMad
A
B
C
UASp-LifeAct-GFP / + ; ; hs-bam / nosGAL4VP16
GFP pMad DAPI
GFP
E-cad
pMad
GFP Ecad DAPI
D
E
UASp-LifeAct-GFP / + ; ; hs-bam / nosGAL4VP16
GFP αSpectrin DAPI
GFP
αSpectrin
Bam
pMad
F
G
H
I
Control
hs-bam
UASp-LifeAct-GFP / + ; ; hs-bam / nosGAL4VP16

**Figure S1, related to Figure 2: GC differentiation phenotypes**

(A-C) Germaria carrying *LifeAct-GFP* driven by *nosGAL4VP16* and a *hs-bam* transgene 3 dpHS immunostained with anti-GFP (green and inset), anti-αSpectrin (magenta and inset), anti-pMad (single channel) and DAPI (blue). Germaria have (A) 16cc with no cytonemes, (B) 8cc with no cytonemes, (C) 4cc with small cytonemes and weak pMad signal. Magenta arrowheads indicate pMad, cyan arrowheads indicate cytonemes. (*) Niche cells, including extra somatic cells in (A) and (B). Scale bar, 5μm.

(D-E) Germaria as in (A), but immunostained with anti-GFP (green and inset), anti-pMad (magenta and inset), anti-Ecad (inset) and DAPI (blue) on 4 dpHS. Germaria show (D) 4cc with small cytonemes and no pMad signal, (E) pMad-positive GSC with large cytonemes. Final panel shows 2x magnification. Magenta arrowheads indicate pMad, cyan arrowheads indicate cytonemes. (*) Niche cells. Scale bar, 5μm.

(F-I) Germaria carrying *LifeAct-GFP* driven by *nosGAL4VP16* (F) and a *hs-bam* transgene (G-I) 3 dpHS, immunostained with anti-GFP (green and inset), anti-pMad (inset), anti-Bam (inset), anti-αSpectrin (grey and inset) and DAPI (blue). Germaria have (F) pMad-positive GSC with cytonemes, (G) 8cc with no cytonemes and extra niche cells, (H) pMad-positive 4cc with small cytonemes and (I) pMad-positive GSC with cytonemes. Magenta arrowheads indicate pMad, cyan arrowheads indicate cytonemes. (*) Niche cells, including extra somatic cells in (G). Scale bar, 5μm.

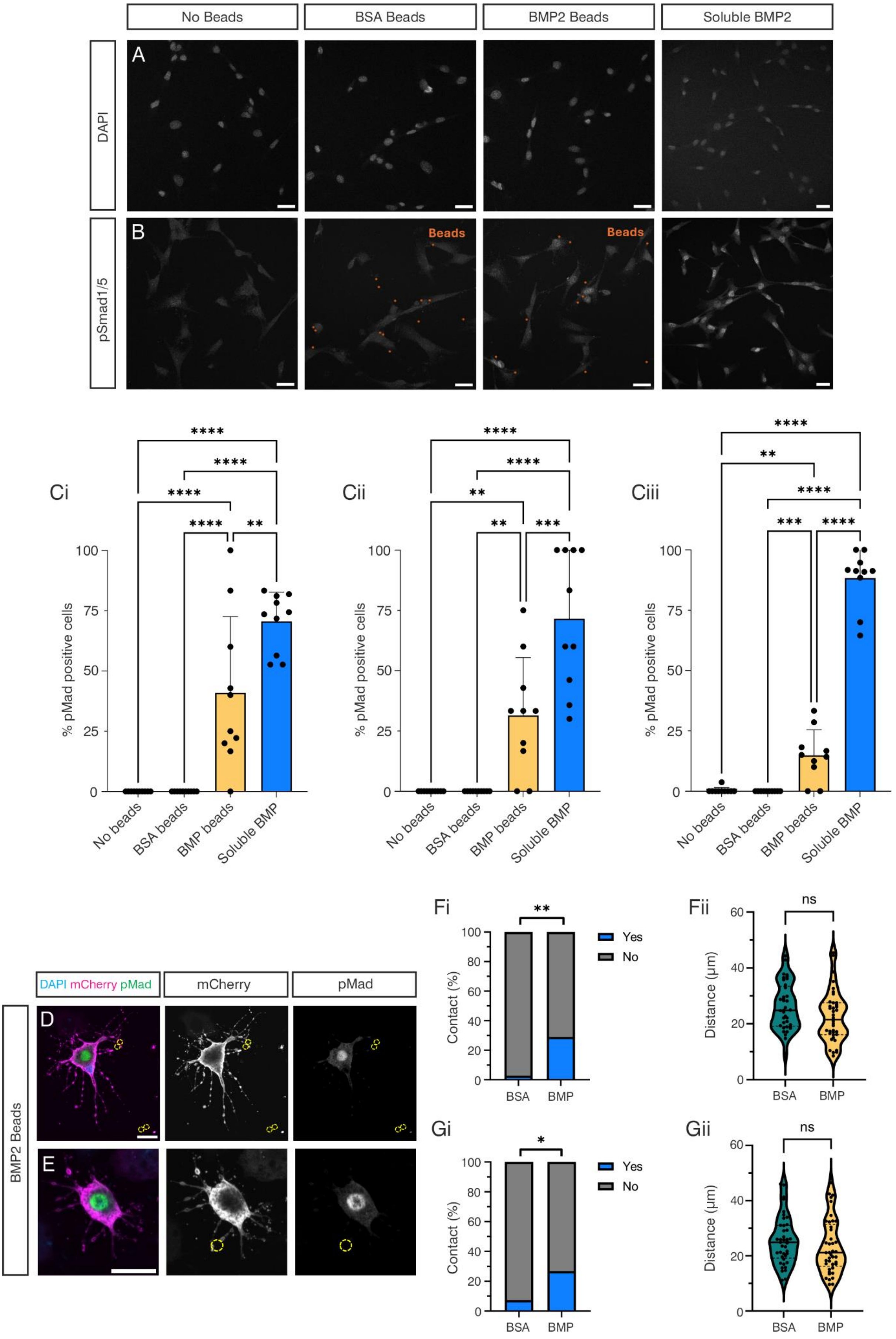
No Beads
BSA Beads
BMP2 Beads
Soluble BMP2
DAPI
pSmad1/5
A
B
Beads
Ci
Cii
Ciii
% pMad positive cells
No beads
BSA beads
BMP beads
Soluble BMP
****
**
***
DAPI mCherry pMad
mCherry
pMad
BMP2 Beads
D
E
Fi
Fii
Gi
Gii
Contact (%)
Yes
No
BSA
BMP
Distance (µm)
ns

**Figure S2, related to Figure 2: Tkv-coated S2R+ cell cytonemes contact BMP beads**

(A, B) C2C12 cells incubated with either BSA coated beads, BMP2 coated beads or soluble recombinant BMP2, immunostained with DAPI (A) and pSmad1/5 (B). Bead locations are marked in orange. Scale bar; 30μm.

(Ci-iii) Quantification of pMad-positive C2C12 cells incubated with no beads, BSA beads, BMP2 beads or soluble BMP2, for 3 biological replicates (replicates 1-3, ordered i-iii). Data analysed using one-way ANOVA with Tukey's multiple comparisons test, n = 10 fields of cells, with on average 20 cells in each. ** p = <0.01, *** p = <0.001, **** p = <0.0001. Data shown as means ± SD.

(D, E) S2R+ cells transfected with Tkv-mCherry and FLAG-Mad expression plasmids were incubated with BMP2 coated beads and immunostained with anti-tdTomato (magenta and inset) and anti-pMad (green and inset). Bead locations are marked with yellow dashed lines. Scale bars 10μm.

(F) (i) Quantification of contacts between BSA or BMP2 coated beads and S2R+ cells. n = 37 (BSA), 38 (BMP2), Fisher's exact test, p = 0.0031. (ii) Quantification of the distance between the bead and the centre of the cell nucleus. Unpaired t test p = 0.0724. Data are for repeat 2.

(Gi) as in (Fi) but for biological repeat 3. n = 40 (BSA), 41 (BMP2), Fisher's exact test, p = 0.0372. (Gii) as in (Fii) but for biological repeat 3. Mann-Whitney test p = 0.1866.

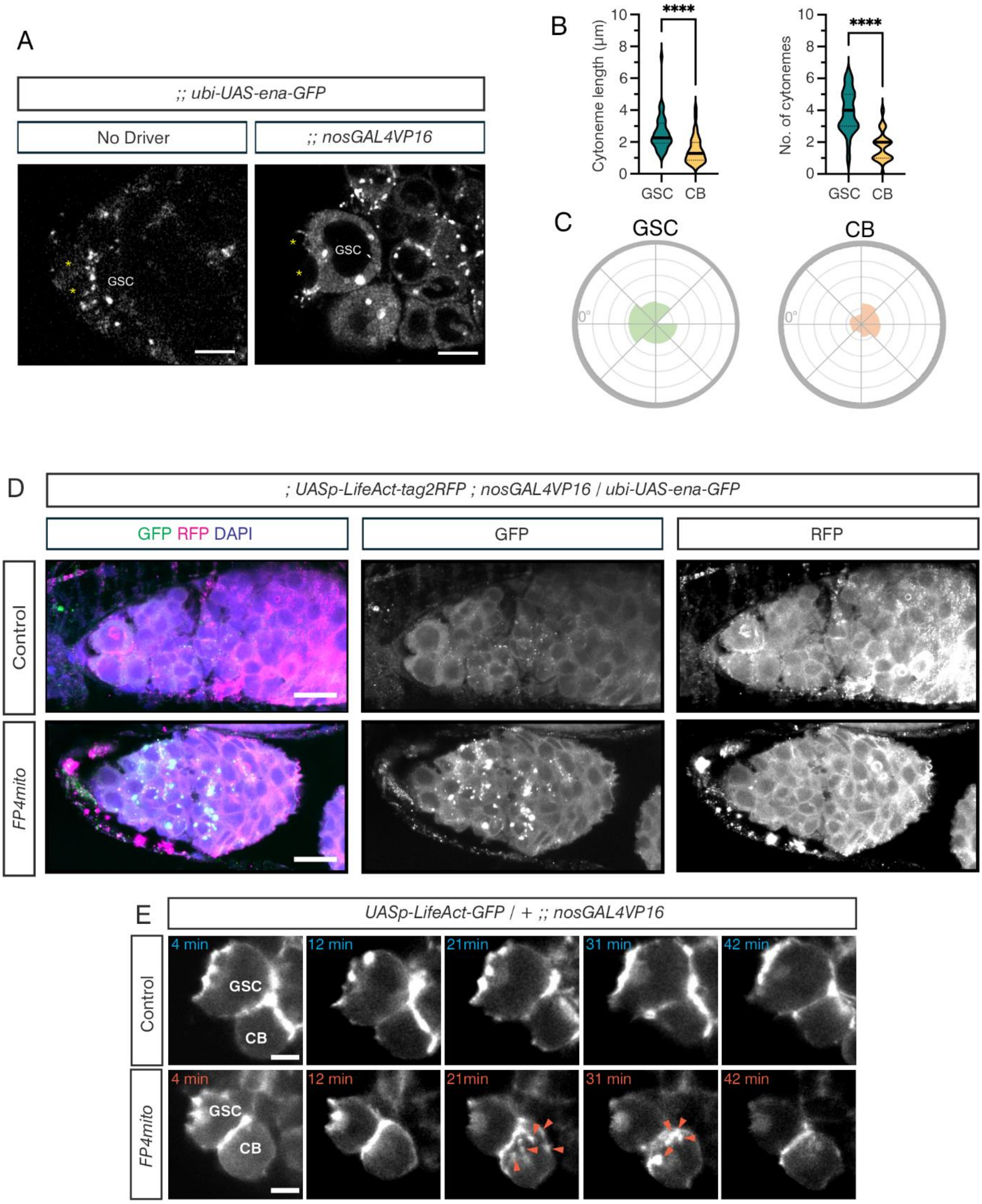
A
;; ubi-UAS-ena-GFP
No Driver
;; nosGAL4VP16
GSC
GSC
B
Cytoneme length (µm)
No. of cytonemes
****
****
GSC CB
GSC CB
C
GSC
CB
0°
0°
D
; UASp-LifeAct-tag2RFP ; nosGAL4VP16 / ubi-UAS-ena-GFP
GFP RFP DAPI
GFP
RFP
Control
FP4mito
E
UASp-LifeAct-GFP / + ;; nosGAL4VP16
Control
FP4mito
4 min
12 min
21min
31 min
42 min
GSC
CB

**Figure S3 related to Figure 3: Ena localisation in *FP4mito* expressing germaria**

(A) GFP fluorescence in live germaria carrying a *ubi-UAS-Ena-GFP* transgene and no driver (left panel) or the *nosGAL4VP16* driver (right panel). A GSC and Cap Cells (*) are labelled. Scale bar, 5µm.

(B, C) Quantitation of (B) cytoneme number (n = 22), length (n = 39 (GSC) and n = 37 (CB) and (C) orientation to the niche (0°), each interval is 10%, in GSCs and CBs (n = 17 GSCs and CBs). Analysis is from live images of *ex vivo* germaria expressing *pUASp-GAP43-Venus* driven by *nosGAL4VP16* (an example image is shown in Figure 3F). Mann-Whitney test, $p = < 0.0001$, $p = < 0.0001$ respectively.

(D) *FP4mito* or control germaria expressing *LifeAct-tag2RFP* and *ubi-UAS-ena-GFP* driven by *nosGAL4VP16* immunostained with anti-RFP (magenta and single channel), anti-GFP (green and single channel) and DAPI (blue). Scale bar, 10µm.

(E) Stills from a live imaging movie of control or *FP4mito* germaria carrying *LifeAct-GFP* driven by *nosGAL4VP16*. Orange arrowheads indicate actin localised intracellularly. Scale bar, 5µm.

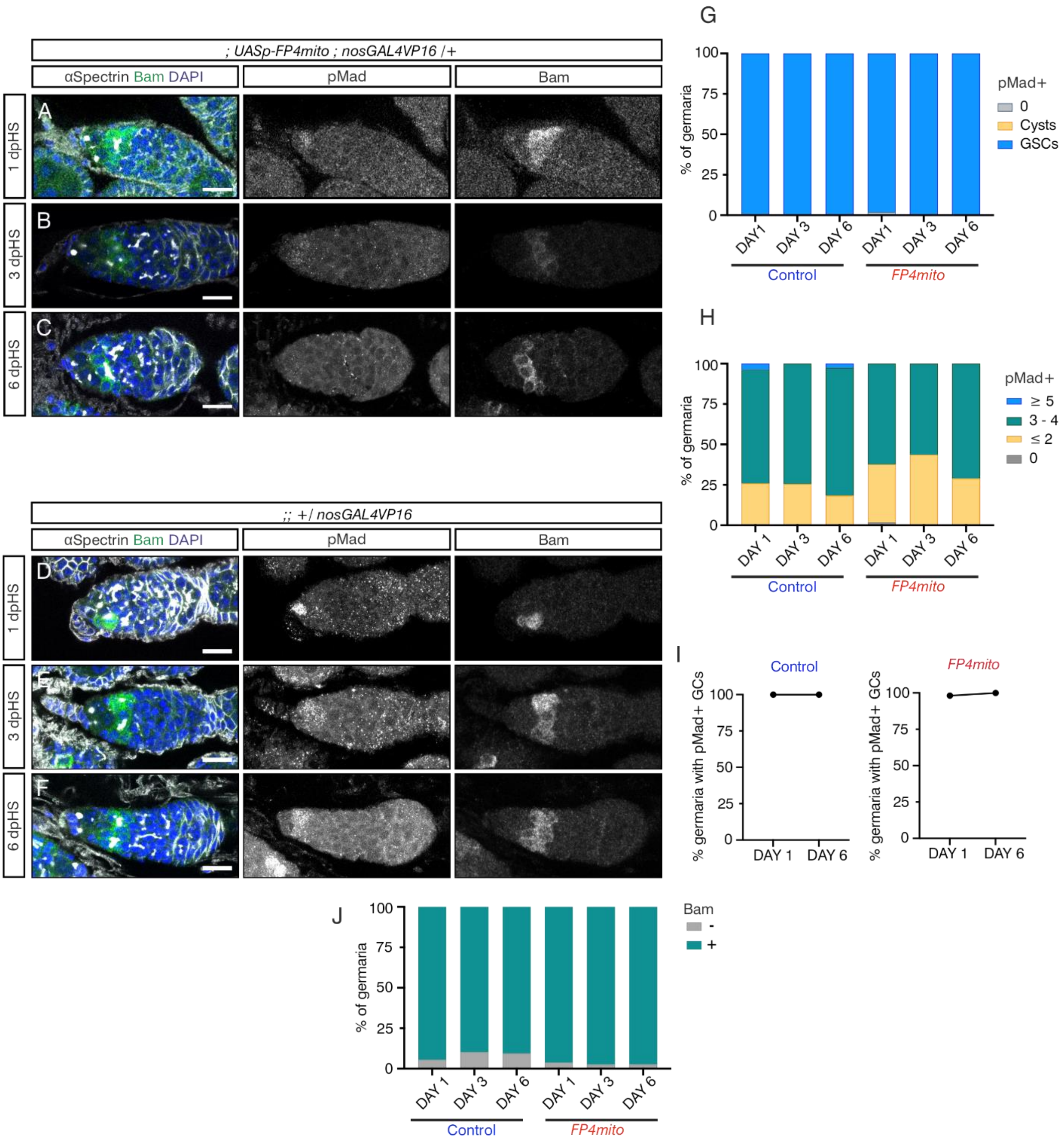


**Figure S4, related to Figure 6: HS does not affect GSC maintenance or differentiation**

(A-C) Control germaria carrying *hs-bam*, *nosGAL4VP16* and *UASp-FP4mito* transgenes immunostained with anti-Bam (green and inset), pMad (inset), anti-αSpectrin (grey) and DAPI (blue). Representative germaria are shown for 1 (A), 3 (B) and 6 (C) dpHS. Scale bar; 10µm.

(D-F) As in (A-C), except the germaria only carry *hs-bam* and *nosGAL4VP16* insertions.

(G-J) Quantification of germaria containing pMad-positive GSCs and cysts (G), total pMad-positive GCs (H), and Bam-positive GCs (J) on 1, 3 and 6 dpHS for control and *FP4mito* shown in (A-F). (I) Comparison of the proportion of pMad-positive cells on day 1 and 6 pHS. n>30 per genotype.

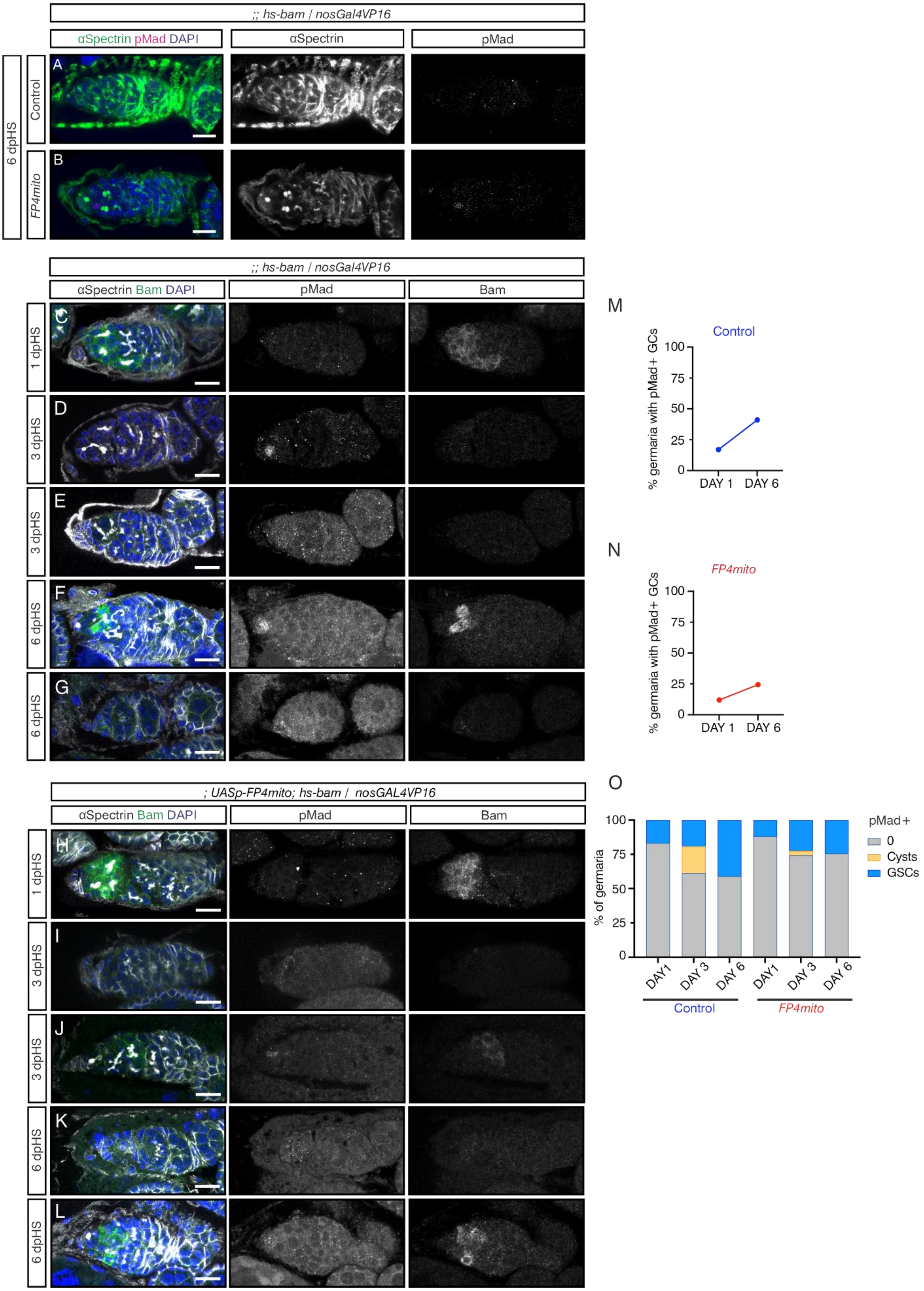
;; hs-bam / nosGal4VP16
αSpectrin pMad DAPI
αSpectrin
pMad
6 dpHS
Control
FP4mito
A
B
;; hs-bam / nosGal4VP16
αSpectrin Bam DAPI
pMad
Bam
C
D
E
F
G
1 dpHS
3 dpHS
3 dpHS
6 dpHS
6 dpHS
; UASp-FP4mito; hs-bam / nosGAL4VP16
αSpectrin Bam DAPI
pMad
Bam
H
I
J
K
L
1 dpHS
3 dpHS
3 dpHS
6 dpHS
6 dpHS
M
Control
% germaria with pMad+ GCs
100
75
50
25
0
DAY 1
DAY 6
N
FP4mito
% germaria with pMad+ GCs
100
75
50
25
0
DAY 1
DAY 6
O
% of germaria
100
75
50
25
0
DAY1
DAY 3
DAY 6
DAY1
DAY 3
DAY 6
Control
FP4mito
pMad+
0
Cysts
GSCs

**Figure S5, related to Figure 6: Ena mislocalisation impairs GSC dedifferentiation**

(A, B) Control (A) and *FP4mito* (B) germaria carrying a *hs-bam* transgene and *nosGAL4VP16* immunostained with anti-pMad (magenta and inset), anti-αSpectrin (green and inset) and DAPI (blue). Representative germaria (other examples are in Figure 6A-D) are shown for 6dpHS. Scale bar; 10μm.

(C-G) Germaria carrying a *hs-bam* transgene and *nosGAL4VP16* immunostained with anti-pMad (inset), anti-αSpectrin (grey), anti-Bam (green and inset) and DAPI (blue). Scale bar; 10μm. Germaria are 1 (C), 3 (D, E) and 6 (F, G) dpHS. (C) 1dpHS shows loss of GSCs indicated by loss of spectrosomes and pMad. GCs adjacent to the niche are Bam-positive. (D) 3dpHS some GCs closest to the niche have become pMad-positive, while others (E) have only pMad-negative cysts. (F) 6 dpHS, some germaria have regained GSCs in the niche, with weak pMad signal. Bam can be detected in the early cysts. (G) Other germaria have lost all early GCs with no pMad or Bam signal detectable.

(H-L) As in (C-G), except germaria also carry the *UASp-FP4mito* transgene. (H) GSCs are lost 1dpHS as indicated by an absence of spectrosomes and pMad signal, and the presence of Bam-positive GCs adjacent to the niche. (I) At 3dpHS, some germaria only have pMad-negative cysts, while others (J) have weakly pMad-positive GCs. (K) At 6 dpHS, some germaria have no pMad or Bam signal, indicative of loss of all early GCs. (L) Other germaria have pMad-positive GSCs reoccupying the niche and Bam-positive early cysts.

(M, N) Comparison of the proportion of pMad-positive cells at day 1 and 6 pHS, for control (M) and *FP4mito* (N). n>30 per genotype.

(O) Quantification of *hs-bam* germaria containing pMad-positive GSCs and cysts on days 1, 3 and 6 pHS for control and *FP4mito* shown in (A-J). n>30 per genotype.

**Table S1.** ***Drosophila*** **stocks used in this study.**

*Drosophila* stocks used are listed with their source.

| | | |
|---|---|---|
| *D. melanogaster; y[1] w[67c23]* | Bloomington | Cat# 6599; RRID:BDSC_6599 |
| *D. melanogaster*; *w*;*<br>*GAL4::VP16-nos* | Bloomington | Cat# 4937; RRID: BDSC_4937 |
| *D. melanogaster;*<br>*M{w[+mC]=UASp-LifeAct.mGFP6}ZH-2A,*<br>*w[*]* | Bloomington | Cat# 58717; RRID: BDSC_58717 |
| *D. melanogaster; w[*]; M{w[+mC]=UASp-Lifeact.TagRFP.H}ZH-51D/CyO* | Bloomington | Cat# 58717; RRID: BDSC_58717 |
| *D. melanogaster;* w[1118]; P{w[+mC]=hs-bam.O}11d/TM3, Sb[1] | Bloomington | Cat# 24637; RRID:BDSC_24637 |
| *D. melanogaster;* P{w[+mC]=hs-bam.O}18d, w[1118] | Bloomington | Cat# 24636; RRID:BDSC_24636 |
| *D. melanogaster; y[1] w[*]; P{w[+mC]=UAS-FLAG-HA-FP4mito}3* | Bloomington | Cat# 58481; RRID:BDSC_58481 |
| *D. melanogaster; w*;P{w[+mC]=UASp-halo7-FP4mito}attP40* | This study | |
| *D. melanogaster; w*;*<br>*bam-GFP* | D. McKearin | (4) |
| *D. melanogaster; w[*]; P{w[+mC]=Ubi-GFP.ena}3* | Bloomington | Cat# 28798; RRID:BDSC_28798 |
| *D. melanogaster; y w M(eGFP, vas-int, dmRFP)ZH-2A; P{CaryP}attP40* | University of Cambridge | Stock 13-20 |
| *D. melanogaster; w[*]; P{w[+mC]=UASp-Venus.GAP43}10* | Bloomington | Cat# 30896; RRID: BDSC_30896 |

**Table S2. Primers use in this study.**

Primer sequences are listed.

| | |
|---|---|
| mito F | CGGGGATCAGATCCGCGGCCGCCAAAATGGCAGAAATCGGTACTGGC |
| mito R | TGAGTCCGGAACTAGTCTCGAGACTAGTAGATCTG |
| Halo7 F | CGAGACTAGTTCCGGACTCAGATCTATCGTAGCT |
| Halo7 R | ACGTTCGAGGTCGACTCTAGAAGCTAGCCTAGGCTCGAGA |

**Table S3. Antibodies used in this study.**

Antibodies used are listed with their source.

| | | | |
|---|---|---|---|
| Anti-GFP (Goat) | 1:500 (IF)<br>1:250<br>(Cytoneme IF) | Abcam | Cat#Ab6673<br>RRID:AB_305643 |
| Anti-pSmad3 Phospho (pS423/425) [EP823Y] (rabbit), recognises pMad | 1:500 (IF, *in vivo*, S2R+)<br>1:250<br>(Cytoneme IF) | Abcam | Cat Ab52903<br>RRID:AB_765068 |
| Anti-Spectin 3A9 (323 or M10-2)<br>(mouse) | 1:50 (IF)<br>1:25 (Cytoneme IF) | Hybridoma Bank | AB_528473<br>RRID:AB_528473 |
| Anti-Enabled 5G2 (mouse) | 1:25 | Hybridoma Bank | AB_528220<br>RRID:AB_528220 |
| Anti-DCAD2 (rat) | 1:25 | Hybridoma Bank | AB_528120<br>RRID:AB_528120 |
| Anti-Vasa (d-260) (rabbit) | 1:500 | Santa Cruz (discontinued) | Cat# sc-30210<br>RRID:AB_793874 |
| Anti-Bam (mouse) | 1:50 | Hybridoma Bank | Cat# bam,<br>RRID:AB_10570327 |
| Anti-GFP (rabbit) | 1:500 (IF)<br>1:250<br>(Cytoneme IF) | Abcam | Cat# ab6556,<br>RRID:AB_305564 |
| Anti-GFP (chicken) | 1:500 (IF)<br>1:250<br>(Cytoneme IF) | Abcam | Cat# ab13970,<br>RRID:AB_300798 |
| Anti-tdTomato (goat) | 1:250 | OriGene | Cat# AB8181-200,<br>RRID:AB_3206272 |
| Anti-Phospho-Smad1 (Ser463/465)/ Smad5 (Ser463/465)/ Smad9 (Ser465/467) (D5B10) (rabbit) | 1:500 (IF, C2C12 cells) | Cell Signaling Technology | Cat# 13820,<br>RRID:AB_2493181 |
| Donkey anti-Rabbit IgG (H+L) Highly Cross-Absorbed Secondary | 1:1000 | Thermo Fisher Scientific | Cat# A-31573<br>RRID:AB_2536183 |

| | | | |
|---|---|---|---|
| Antibody, Alexa Fluor™ 647 | | | |
| Donkey anti-Goat IgG (H+L) Cross-Absorbed Secondary Antibody, Alexa Fluor™ 488 | 1:1000 | Thermo Fisher Scientific | Cat# A-11055 RRID:AB_2534102 |
| Donkey anti-Rat IgG (H+L) Highly Cross-Absorbed Secondary Antibody, Alexa Fluor™ 647 | 1:1000 | Thermo Fisher Scientific | Cat# A78947, RRID:AB_2910635 |
| Donkey anti-Rat IgG (H+L) Highly Cross-Absorbed Secondary Antibody, Alexa Fluor™ 594 | 1:1000 | Thermo Fisher Scientific | Cat# A-21209, RRID:AB_2535795 |
| Donkey anti-Rabbit IgG (H+L) Highly Cross-Absorbed Secondary Antibody, Alexa Fluor™ 594 | 1:1000 | Thermo Fisher Scientific | Cat# A-21207, RRID:AB_141637 |
| Donkey anti-Mouse IgG (H+L) Highly Cross-Absorbed Secondary Antibody, Alexa Fluor™ 594 | 1:1000 | Thermo Fisher Scientific | Cat# A-21203, RRID:AB_2535789 |
| Donkey anti-Mouse IgG (H+L) Highly Cross-Absorbed Secondary Antibody, Alexa Fluor™ 1:1000555 | 1:1000 | Thermo Fisher Scientific | Cat# A-31570, RRID:AB_2536180 |
| Donkey anti-Rabbit IgG (H+L) Highly Cross-Absorbed Secondary Antibody, Alexa Fluor™ 488 | 1:1000 | Thermo Fisher Scientific | Cat# A-21206, RRID:AB_2535792 |

| | | | |
|---|---|---|---|
| Donkey anti-Mouse IgG (H+L) Highly Cross-Absorbed Secondary Antibody, Alexa Fluor™ 647 | 1:1000 | Thermo Fisher Scientific | Cat# A-31571, RRID:AB_162542 |
| Goat anti-Chicken IgY (H+L) Secondary Antibody, Alexa Fluor™ 488 | 1:1000 | Thermo Fisher Scientific | Cat# A-11039, RRID:AB_2534096 |
| Donkey anti-Rabbit IgG (H+L) Highly Cross-Absorbed Secondary Antibody, Alexa Fluor™ 790 | 1:1000 | Thermo Fisher Scientific | Cat# A11374, RRID:AB_2534145 |
| Donkey anti-Goat IgG (H+L) Cross-Absorbed Secondary Antibody, Alexa Fluor™ 594 | 1:1000 | Thermo Fisher Scientific | Cat# A-11058, RRID:AB_2534105 |
| Zenon™ Alexa Fluor™ 555 Mouse IgG1 Labeling Kit | NA | Thermo Fisher Scientific | Cat# Z25005, RRID:AB_2736948 |

**SI References**

1. B. Sutcliffe *et al.*, Second-Generation Drosophila Chemical Tags: Sensitivity, Versatility, and Speed. *Genetics* **205**, 1399-1408 (2017).
2. S. G. Wilcockson, H. L. Ashe, Drosophila Ovarian Germline Stem Cell Cytocensor Projections Dynamically Receive and Attenuate BMP Signaling. *Dev Cell* **50**, 296-312 e295 (2019).
3. J. Schindelin *et al.*, Fiji: an open-source platform for biological-image analysis. *Nat Methods* **9**, 676-682 (2012).
4. D. Chen, D. McKearin, Dpp signaling silences bam transcription directly to establish asymmetric divisions of germline stem cells. *Curr Biol* **13**, 1786-1791 (2003).
